# Repression of Dlx1/2 Signaling by *Nolz-1/Znf503* is Essential for Parcellation of the Striatal Complex into Dorsal and Ventral Striatum

**DOI:** 10.1101/463398

**Authors:** Kuan-Ming Lu, Shih-Yun Chen, Hsin-An Ko, Ting-Hao Huang, Janice Hsin-Jou Hao, Yu-Ting Yan, Sunny Li-Yun Chang, Sylvia Evans, Fu-Chin Liu

## Abstract

The division of the striatum into dorsal and ventral districts is of central clinical importance. The dorsal striatum is differentially affected in Huntington’s disease, dopamine in the ventral striatum is differentially spared in Parkinson’s disease, and human brain imaging studies implicate the ventral striatum in addictive disorders. If fits that the dorsal striatum contains the cells of origin of the direct and indirect basal ganglia pathways for motor control. The ventral striatum is a node in neural circuits related to motivation and affect. Despite these striking neurobiologic contrasts, there is almost no information about how the dorsal and ventral divisions of the striatum are set up during development. Here, we demonstrate that interactions between the two key transcription factors Nolz-1 and Dlx1/2 control the migratory paths of developing striatal neurons to the dorsal or ventral striatum. Moreover, these same transcription factors control the cell identity of striatal projection neurons in both the dorsal and ventral striatum including the cell origin of the direct and indirect pathways. We show that Nolz-1 suppresses Dlx1/2 expression. Deletion of Nolz-1 or over-expression of Dlx1/2 can produce a striatal phenotype characterized by withered dorsal striatum and a swollen ventral striatum, and that we can rescue this phenotype by manipulating the interactions between Nolz-1 and Dlx1/2 transcription factors. This evidence suggests that the fundamental basis for divisions of the striatum known to be differentially vulnerable at maturity is already encoded by the time embryonic striatal neurons begin their migrations into the developing striatum.

## INTRODUCTION

The striatum of the basal ganglia plays diverse roles in the regulation of movement, reward, motivation and cognitive behaviors, and dysfunction of the striatum is involved in a number of neurological and psychiatric disorders (Crittenden and Graybiel, 2011; Gerfen and Surmeier, 2011; Wichmann and DeLong, 1996). The diverse neurological functions stem from the fact that the striatum is a complex structure. The striatum is a two-tier system that comprises cytoarchitecturally similar but functionally distinct dorsal and ventral striatum. The dorsal striatum consists of the caudate-putamen (CP) that controls movement and habit learning (Graybiel, 2008). The ventral striatum consists of the nucleus of accumbens (NAc) and olfactory tubercle (OT) that regulates motivation, reward learning and drug addiction (Grueter et al., 2012; Heimer et al., 1995; Russo et al., 2010). Despite the extensive knowledge of the structure and function of the striatal complex in adulthood, little is yet known about how the dorsal and ventral striatum are differentially specified during development.

Because CP, NAc and OT neurons express similar profiles of transcription factors and neurotransmission-related molecules, and exhibit cellular morphology similar to that of medium-sized spiny neurons (MSNs) (Gerfen and Wilson, 1996; Heimer et al., 1995), they may share developmental origins in neurogenesis. This is supported by homotopic transplantation studies, which show that donor-derived cells grafted from the lateral ganglionic eminence (LGE, striatal anlage) are distributed throughout CP, NAc and OT neurons of host brains (Wichterle et al., 2001). The LGE is divided into dorsal and ventral LGE (Stenman et al., 2003). The dorsal and ventral LGE give rise to interneurons in the olfactory bulb and projection neurons in the CP, respectively. It is yet unclear whether the progenitors of CP, NAc and OT neurons are localized in specific domains within the LGE, and/or they are derived from temporal progression of progenitors that differentiate at different time windows, through combinatorial expression of transcription factor codes delineated in progenitor domains of the LGE (Flames et al., 2007).

In an attempt to decipher mechanisms underlying the developmental construction of dorsal and ventral striatum, we performed genomewide comparisons of gene expression patterns of dorsal and ventral parts of the LGE. We identified a number of genes that were differentially expressed in the dorsal and ventral developing striatum. We focused on *Nolz-1*/*Znf503/Zfp503* that was expressed at high levels in developing dorsal striatum. Here, we report that *Nolz-1* plays a novel role in the regulation of cell type specification and neuronal migration in the dorsal and ventral striatum during development. *Nolz-1* null mutation not only resulted in aberrant differentiation of striatal neurons of the dorsal and ventral striatum, but also induced abnormal enlargement of ventral striatum at the expense of dorsal striatum. The distorted striatal complex in the *Nolz-1* mutant brain was primarily caused by an abnormal Dlx1/2-dependent cell migration, which drove aberrant migration of striatal cells from the dorsal toward the ventral striatum. Therefore, repression of Dlx1/2 signaling by Nolz-1 is required to restrict migration of striatal cells to specific sub-domains, which leads to parcellation of the striatal complex into the dorsal and ventral striatum.

## RESULTS

### Identification of genes differentially enriched in the dorsal or ventral striatum during development

To search for genes that are differentially expressed in developing dorsal and ventral striatum, we dissected dorsal and ventral parts of the lateral ganglionic eminence (LGE, striatal anlage) in E13.5 mouse forebrain (Fig. 1A), and then performed genomewide microarray analysis. We compared gene expression profiles between dorsal and ventral parts of the LGE. Ingenuity Pathway Analysis revealed differentially expressed genes in several categories, including Sonic Hedgehog Signaling, FGF Signaling, TGF-β Signaling, Human Embryonic Stem Cell Pluripotency, Transcriptional Regulatory Network in Embryonic Stem Cells, Axonal Guidance Signaling and others, and these genes included growth factors, and transcription factors (Supplemental Table S1). qRT-PCR assays confirmed dorsal-enriched genes *Wnt7b* and *Nolz-1/Znf503*, and ventral-enriched genes *Acvr2a, fgf14, fgf15, Otx2, Gli1, Zic1, Nrp2, Plxnc1, Nts* and *Calb1* (Fig. 1A-B).

**Figure 1.**
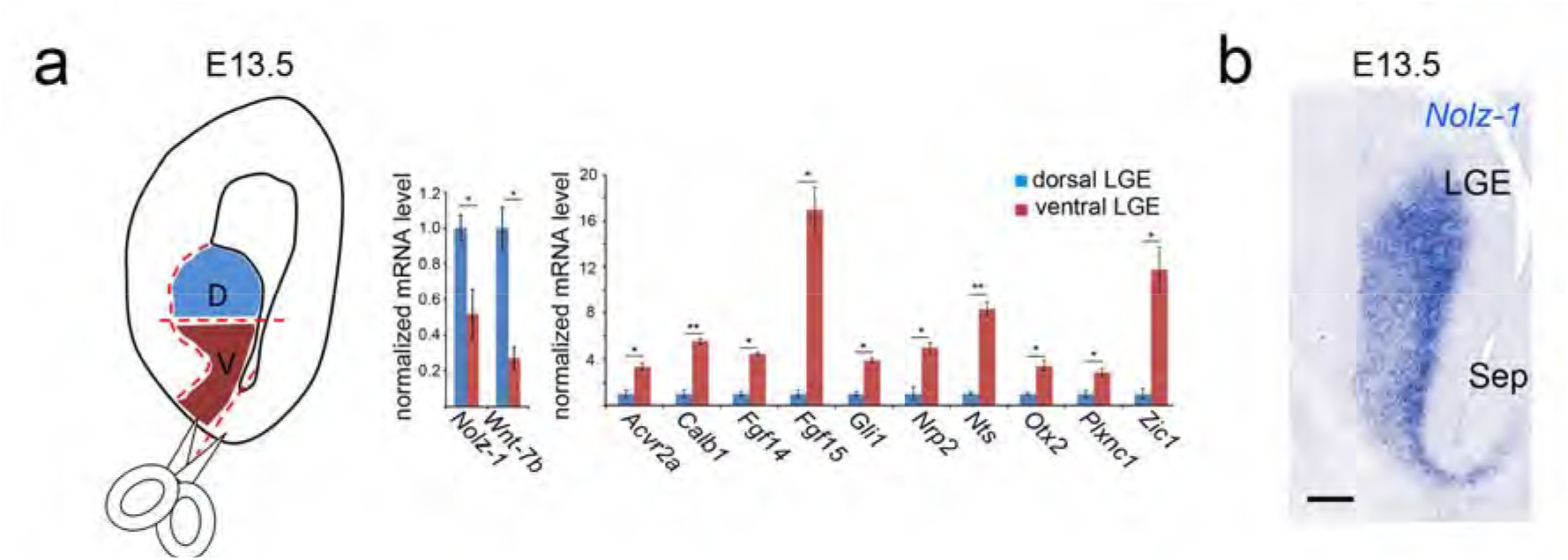
Identification of dorsal striatum- or ventral striatum-enriched genes in developing mouse forebrain. **A:** Schematic drawing of dissection of dorsal or ventral parts of the lateral ganglionic eminence (LGE, striatal anlage) in E13.5 mouse forebrain. qRT-PCR assays of E13.5 LGE show that *Nolz-1* and *Wnt-7b* are expressed at higher levels in the dorsal than ventral part of LGE, while *Acvr2a, Calb1, Fgf14, Fgf15, Gli1, Nrp2, Nts, Otx2, Plxnc1* and *Zic1* are expressed at higher levels in the ventral than dorsal part of LGE. **B:** *Nolz-1* mRNA is expressed at high levels in the dorsal part of E13.5 LGE. *, p < 0.05; **, p < 0.01, n =3/group. Scale bar, 100 μm in B.

### *Nolz-1* regulates differentiation and axonal outgrowth/navigation of striatonigral neurons

Among genes enriched in the dorsal striatum of E13.5 mouse brain, *Nolz-1* was of particular interest (Fig. 1B). We have previously shown that *Nolz-1* is a developmentally regulated striatum-enriched gene in rat brain (Chang et al., 2004). We generated floxed *Nolz-1^fl/fl^* mice for studying *Nolz-1* function (Supplemental Fig. S1). *Nolz-1^fl/fl^* mice were intercrossed with Protamine-Cre mice to generate germline *Nolz-1*^-/-^ knockout (KO) mice. *Nolz-1*^-/-^ KO mice died immediately after birth. Immunostaining of Nolz-1 confirmed the absence of Nolz-1 protein in E18.5 *Nolz-1* KO striatum (Supplemental Fig. S1).

We first examined whether general differentiation of striatal neurons was affected in *Nolz-1* KO brains by immunostaining with striatal differentiation markers, including Foxp2, DARPP-32 and CalDAG-GEFI (Kawasaki et al., 1998; Ouimet et al., 1984; Takahashi et al., 2003). Foxp2 was significantly reduced in *Nolz-1* KO striatum at E18.5 (Fig. 2A). DARPP-32 and CalDAG-GEFI, markers of developing striosomal and matrix compartments, respectively, were also markedly decreased in KO striatum (Fig. 2B, C).

**Figure 2.**
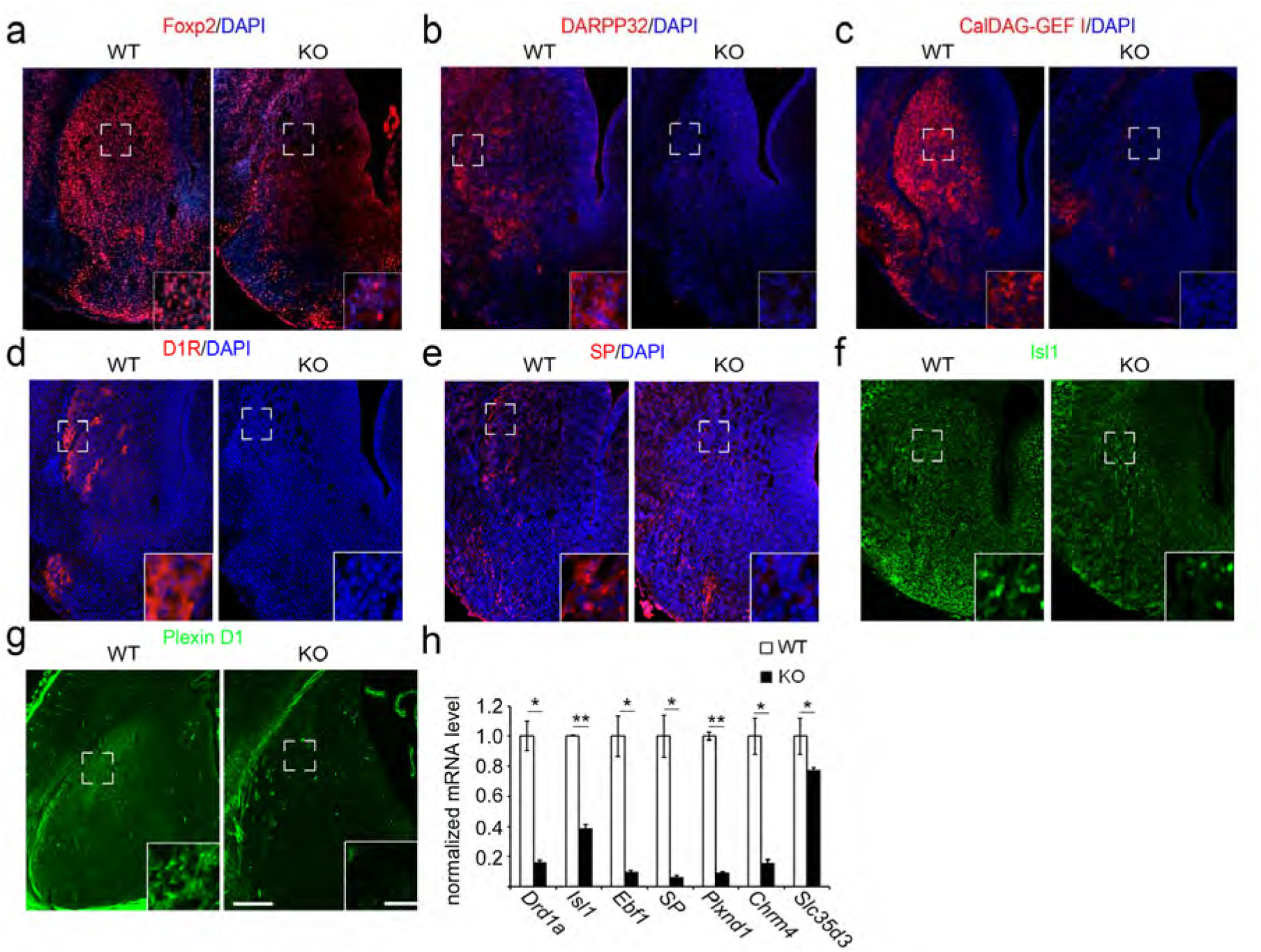
Striatonigral-enriched gene expression is decreased in the *Nolz-1* knockout striatum. **A:** Foxp2 is decreased is decreased in *Nolz-1* knockout (KO) striatum compared to wild-type (WT) striatum of E18.5 brains. **B, C:** DARPP-32-positive striosomes (B) and CalDAG-GEF I-positive matrix (C) are decreased in *Nolz-1* KO striatum. **D-F:** Dopamine D1 receptor (D1R, D), substance P (SP, E), Isl1 (F) and Plexin D1 (G), markers of striatonigral neurons, are decreased in *Nolz-1* KO striatum. The bracketed regions are shown at high magnification in insets. **H:** qRT-PCR assay shows significant reductions of striatonigral-enriched genes in E18.5 *Nolz-1* KO striatum. *, p < 0.05; **, p < 0.01, n =3/group. Scale bars, 200 μm in G for A-G, 30 μm in inset.

Striatal projection neurons consist of striatonigral and striatopallidal neurons (Gerfen and Wilson, 1996). We further asked whether *Nolz-1* mutation selectively affected subtypes of striatal projection neurons. Immunostaining showed significant reductions in Dopamine D1 receptor (D1R), Substance P (SP), Plexin D1 (PlxnD1) and Isl1, markers of striatonigral neurons (Ehrman et al., 2013; Lobo et al., 2006; Lu et al., 2014), in E18.5 *Nolz-1* KO striatum (Fig. 2D-G). Consistently, qRT-PCR analysis indicated that a repertoire of mRNAs encoding striatonigral-enriched genes, including *D1R/Drd1a*, *Isl1, Ebf1, SP*, *PlxnD1*, *Chrm4* and *Slc35d3* were also markedly decreased in *Nolz-1* KO striatum (Fig. 2H). DiI anterograde tracing further showed a reduction of DiI-labeled striatonigral axons along the projection pathway, from the diencephalon and telencephalon boundary through the cerebral peduncles to the substantia nigra of the midbrain (Supplemental Fig. S2A). The defective striatonigral pathway was also evident in Isl1-Cre;Nolz1^fl/fl^;CAG-CAT-EGFP conditional KO brains in which marked reduction of striatonigral GFP^+^ axons was found along the routes of striatonigral axonal projections (see below, Supplemental Fig. S2B).

### Nolz-1 suppresses striatopallidal-enriched genes in developing striatonigral neurons

In dramatic contrast to the reduction of striatonigral-enriched genes, immunostaining showed significant increases of D2R and Met-Enkephalin (Enk), markers of striatopallidal-enriched genes, in E18.5 *Nolz-1* KO striatum (Fig. 3A, B). *In situ* hybridization also revealed a robust increase in *Drd2* mRNA in *Nolz-1* KO striatum throughout rostrocaudal levels (Fig. 3C, D). qRT-PCR assays further confirmed significant increases in mRNAs encoding striatopallidal-enriched genes (Lobo et al., 2006), including *Drd2 (D2R*), *preproenkephalin* (*Penk*) and *adenosine receptor A2a* (*A2aR*) in E18.5 *Nolz-1* KO striatum, although *Gpr6* was decreased in KO striatum (Fig. 3E).

**Figure 3.**
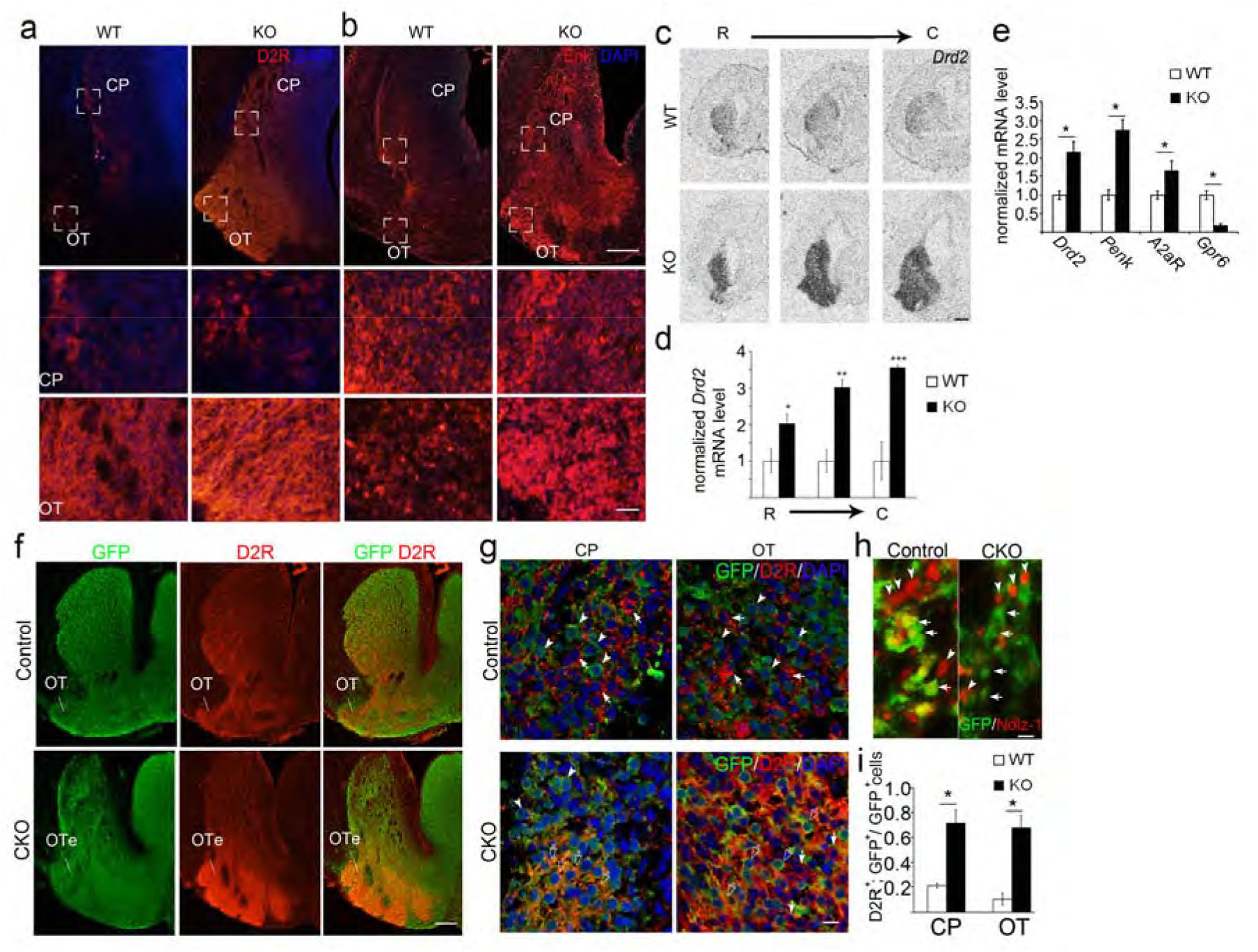
Striatopallidal-enriched genes are de-repressed in striatonigral cells of the *Nolz-1* knockout striatum. **A, B:** Dopamine D2 receptor (D2R) and enkephalin (Enk), markers of striatopallidal neurons, are expressed in both the caudoputamen (CP) and olfactory tubercle (OT) of wild-type (WT) E18.5 brains. D2R and Enk are markedly increased in both CP and OT of *Nolz-1* knockout (KO) brains. The bracketed regions in CP and OT are shown at high magnification in the second and third rows of panels. **C, D:** *In situ* hybridization shows robust de-repression of *Drd2* mRNA in both dorsal and ventral striatum of *Nolz-1* KO brain from rostral (R) to caudal (C) levels. **E:** qRT-PCR assay confirms up-regulation of striatopallidal-enriched genes in E18.5 *Nolz-1* KO striatum except that *Gpr6* is reduced. **F:** Deletion of *Nolz-1* in Isl1 cell lineages results in hyperplasia of ventral striatum and hypoplasia of dorsal striatum in Isl1-Cre;Nolz1^fl/fl^;CAG-CAT-EGFP conditional knockout (CKO) brains compared to control E18.5 Isl1-Cre;Nolz1^fl/+^;CAG-CAT-EGFP brains. **G, I:** D2R (arrows) and GFP (arrowheads) are not co-localized in many striatal cells of CP and OT in control brains. Co-localization of GFP and D2R (empty arrows) are increased in striatal cells of CP and OT-like regions in CKO brains. **H:** Validation of successful deletion of *Nolz-1* in GFP-expressing Isl1 cell lineages in the striatum by the absence of Nolz-1 immunoreactivity (red, arrowheads) in GFP-positive cells (green, arrows) of CKO brains. OTe: enlarged OT. *, p < 0.05; **, p < 0.01; ***, p < 0.001, n =3/group. Scale bars, 200 μm in B, 500 μm in C, 200 μm in F, 10 μm in G, H (B bottom panel, 50 μm).

We further asked whether *Nolz-1* mutation caused de-repression of striatopallidal genes in striatonigral cells by selective deletion of *Nolz-1* in striatonigral cells using Isl1-Cre mice (Ehrman et al., 2013; Lu et al., 2014). Double immunostaining confirmed successful deletion of Nolz-1 protein in GFP-expressing Isl1 cell lineages in Isl1-Cre;Nolz1^fl/fl^;CAG-CAT-EGFP E18.5 mutant brains (Fig. 3I). D2R and GFP were not co-localized in many striatal cells of CP and OT in control Isl1-Cre;Nolz1^fl/+^;CAG-CAT-EGFP brains (Fig. 3F, G). In contrast, co-localization of GFP and D2R was significantly increased in striatal cells of CP and OT-like regions in Isl1-Cre;Nolz1^fl/fl^;CAG-CAT-EGFP mutant brains (CP: WT, 21.2 ± 1.9 % vs. KO 71.7 ± 10.2 %, p < 0.05; OT: WT, 10 ± 3.9 % vs. KO,

69.2 ± 8 %, p < 0.05, n = 3, Fig. 3F, G). These findings suggested that selective deletion of *Nolz-1* in Isl1^+^ striatonigral cell lineages de-repressed striatopallidal genes in striatonigral cells.

### Nolz-1 mutation induces hypoplasia of dorsal striatum but hyperplasia of ventral striatum

Increased D2R and Enk expression marked dramatical structural alteration of the striatal complex in E18.5 *Nolz-1* KO brains. The mutant striatal complex consisted of smaller dorsal striatum but larger ventral striatum when compared to wildtype brains (Fig. 3A, B, C). Similar structural changes were observed in Isl1-Cre;Nolz1^fl/fl^;CAG-CAT-EGFP conditional KO brains (Fig. 3F), suggesting that selective deletion of *Nolz-1* in differentiating striatonigral cells was sufficient to induce hypoplasia and hyperplasia of the dorsal and ventral striatum, respectively. Intriguingly, there were streams of GFP^+^ cells that appeared to extend from the dorsal toward the enlarged ventral mutant striatum (Fig. 3F).

### Altered gene expression patterns in the ventral striatum of Nolz-1 KO brains

The formation of the abnormal ventral striatum was detected as early as E13.5, as shown by the expansion of Meis2 immunostaining into the ventral striatum of *Nolz-1* KO brains (Supplemental Fig. S3A). To characterize the cell types in the mutant ventral striatum, we examined ventral striatum-enriched or dorsal striatum-enriched genes that were identified by our microarray analysis and also from the Allen Brain Atlas. Ventral striatum-enriched genes *Acvr2a* and *Npy1r* were decreased in the ventral striatum of E18.5 *Nolz-1* KO brains (Supplemental Fig. 3B). Dorsal striatum-enriched genes, *Ebf1* and *Astn2*, remained at low levels in the ventral KO striatum, although *Ebf1* and *Astn2* were decreased in dorsal KO striatum (Supplemental Fig. 3B). Expression of Sox1, a gene required for cell migration in the ventral striatum (Ekonomou et al., 2005), was reduced in ventral KO striatum (Supplemental Fig. S3C). Expression of Brn4, which was expressed at high levels in OT and the junction between dorsal and ventral striatum in wildtype brains (Long et al., 2009), was decreased in KO striatum (Supplemental Fig. S3D). Neuropilin 2 (Nrp2) is expressed in OT, but not in CP (Long et al., 2009). In wildtype basal forebrain, a continuous Nrp2^+^ band extended from the septum across OT to the piriform cortex. In contrast, an indented band that comprised weak Nrp2^+^ regions was found in mutant OT-like regions (Supplemental Fig. S3E), suggesting that cells in the expanded mutant OT regions may not differentiate normally or that OT expansion may arise from cells aberrantly immigrating from other regions.

### Cell proliferation is not altered in the germinal zone of the Nolz-1 mutant striatal anlage

Structural changes in *Nolz-1* KO striatal complex might result from altered progenitor populations in striatal anlage. The expression pattern of *Ascl1/Mash1* mRNA, a proneural gene (Ma et al., 1997), was not altered in E13.5 and E16.5 KO striatum (Supplemental Fig. S4). We further examined the proliferation of progenitor cells in LGE by pulse-labeling of proliferating cells with bromodeoxyuridine (BrdU) for 1 hr. The proliferation index (BrdU^+^;Ki67^+^ cells/total Ki67^+^ cells) was not changed in E12.5 LGE and E15.5 striatum of *Nolz-1* KO brains (Supplemental Fig. S5A). We also assayed the cell cycle exit index 24 hr after BrdU injection. The cell cycle exit index (BrdU^+^;Ki67^-^ cells/total BrdU^+^ cells) was not changed in E12.5 LGE and E15.5 striatum of *Nolz-1* KO brains (Supplemental Fig. S5A). Immunostaining of phospho-histone 3 (PH3), a mitotic marker, did not reveal changes in the number of PH3^+^ cells in the ventricular zone (VZ) or subventricular zone (SVZ) of E12.5 LGE. or E15.5 striatum of *Nolz-1* KO brains (Supplemental Fig. S5B).

Previous studies have suggested that the ventral part of the LGE, septoeminential sulcus (SES) and rostromedial telencephalic wall (RMTW), give rise to OT neurons at early embryonic stages (Garcia-Moreno et al., 2008), and that the ventral parts of VZ in the LGE and septum give rise to NAc neurons (Heimer et al., 1997). We found no marked changes in proliferation index (BrdU^+^;Ki67^+^ cells/total Ki67^+^ cells) within any of these progenitor regions in E12.5 and E13.5 *Nolz-1* KO brains (Supplemental Fig. S5C).

### Cell apoptosis is not changed in the Nolz-1 mutant striatum

We then investigated whether abnormal cell death occurred in *Nolz-1*^-/-^ KO striatum by TUNEL assays. No significant changes in TUNEL^+^ apoptotic cells were found in E12.5, E14.5 and E16.5 KO striatum (Supplemental Fig. S6A, B), suggesting that *Nolz-1* is not required for striatal cell survival.

### Abnormal redistribution of striatal cells from the dorsal to the ventral mutant striatum

Since cell proliferation and apoptosis were not changed in *Nolz-1* KO striatum, we next tested whether the concurrent hypoplasia of dorsal striatum and hyperplasia of ventral striatum resulted from redistribution of striatal cells from the dorsal to ventral KO striatum. We pulse labeled early-born and late-born striatal cells with BrdU at E12.5 and E15.5, respectively, and then analyzed the distribution of BrdU-labeled cells at E18.5. Quantitative analysis indicated that the number of BrdU^E12.5^ cells was decreased by 60% in dorsal KO striatum, but was increased by 80% in ventral KO striatum (dorsal and ventral striatum, p < 0.01, n = 3, Supplemental Fig. S7A, C). Similar results were found with late-born BrdU^E15.5^ cells in KO striatum (dorsal: p < 0.05; ventral, p < 0.01, n = 3, Supplemental Fig. S7B, D). As the total numbers of BrdU^E12.5^ cells or BrdU^E15.5^ cells were not markedly changed in the entire KO striatum (p > 0.05, n = 3, Supplemental Fig. S7C, D), these results suggested an abnormal redistribution of striatal cells from dorsal to ventral KO striatum.

### Aberrant migration of striatal cells from dorsal to ventral mutant striatum

The redistribution of BrdU cells implicated abnormal cell migration with an increased number of cells migrating from the dorsal to ventral KO striatum. We then examined this hypothesis by three sets of experiments. Firstly, we focally labeled cells in the dorsal part of the germinal zone of E14.5 LGE with the CFDA cell tracker, and then traced the migration of CFDA-labeled cells 31 hr later *in vivo*. Owing to technical limitations, E14.5 was the earliest time point for which we succeeded for this set of experiments. The successful CFDA injection site was defined as a ~400 μm-radius circle centering at the intensive CFDA-labeled VZ of dorsal LGE (Fig. 5A). CFDA^+^ cells were found to migrate from the dorsomedial toward the ventrolateral parts of wildtype and *Nolz-1* KO LGE. For quantification, we divided the LGE into three equidistant concentric zones (200 μm apart) with the epicenter at the dorsomedial germinal zone where CFDA tracer was deposited, and counted the percentage of CFDA^+^ cells within each zone by dividing CFDA^+^ cells within each zone by the total number of CFDA^+^ cells (Fig. 5A, B). We found that the percentage of CFDA^+^ cells tended to decrease within zone I of *Nolz-1* KO LGE. In contrast, the percentage of CFDA^+^ cells within zone II significantly increased by 76.4% in *Nolz-1* KO relative to control wildtype LGE (zone II, p < 0.05, Fig. 5A, B). These findings indicate that the distribution of CFDA^+^ cells as a whole shifted more ventrolaterally in KO LGE compared to that in wildtype LGE, suggesting that *Nolz-1* null mutation promoted cell migration from dorsal toward ventral striatum.

Secondly, we performed a time course study to trace the migratory routes of cells by pulse labeling striatal cells with BrdU at E12.5, and then chased the cells at 24, 48 and 72 hr following BrdU injection. We plotted the distribution of BrdU^E12.5^ cells, and calculated the percentage of BrdU^E12.5^ cells in each 100 μm equidistant concentric ring. The results showed no changes in BrdU^E12.5^ cells at 24 hr and 48 hr in *Nolz-1* KO brains. By 72 hr, 38.2% cells migrated into the 400-500 μm region, approximately the MZ of the dorsal striatum in wildtype brains. Comparing to wildtype brains, the number of BrdU^E12.5^ cells was decreased by 39.5% in the 400-500 μm region (p < 0.05, n = 3, Fig. 4C, D), but the number of BrdU^E12.5^ cells was increased by 75.4% in the 600-700 μm region where the presumptive mutant OT was located (p < 0.01 n = 3, Fig. 4C, D). As the total numbers of BrdU^E12.5^ cells were not changed in KO brains at 24, 48 and 72 hr compared to wild-type brains (p > 0.05, n = 3, Fig. 4E), these results suggest that by 72 hr, some BrdU^E12.5^ mutant cells do not stop migration, but instead continue migrating to OT-like regions.

**Figure 4.**
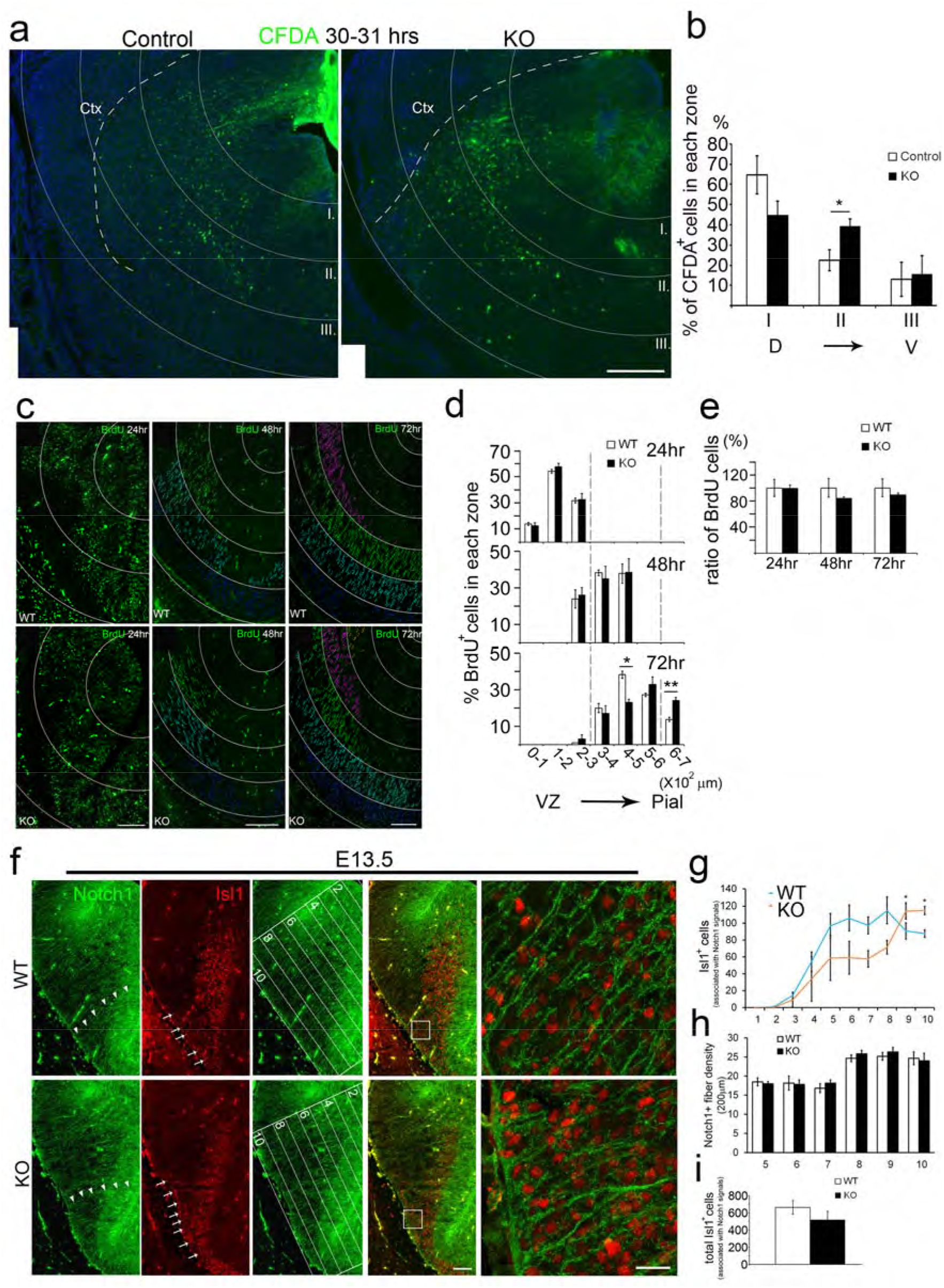
*In vivo* tracing of aberrantly migrating cells in developing striatum of *Nolz-1* knockout brains. **A, B:** CFDA is locally injected into the dorsal part of E14.5 striatum, and the brains are harvested 31 hr later. The striatum is overlaid with dorsal ventricular zone centered-concentric rings with 200 μm apart for cell counting in each zone (zone I to III). Comparing to control brains, more CFDA^+^ cells migrate ventrolaterally into zone II (400-600 μm) in *Nolz-1* knockout (KO) brains. **C:** The migratory pattern of BrdU^E12.5^ cells in the striatum at 24, 48 and 72 hr after BrdU injection. The distribution of BrdU^E12.5^ cells in the striatum are traced in concentric rings with 100 μm apart. **D:** No significant difference of BrdU^E12.5^ cells were found at 24 and 48 hr between wildtype and *Nolz-1* KO brains. By 72 hr, BrdU^E12.5^ cells are decreased in the 400-500 μm region, but increased in the 600-700 μm region in KO brains compared to wildtype brains. **E:** The total number of BrdU^E12.5^ cells at each time point was not different between the wild-type and KO brains. **F:** Double immunostaining of Notch1 and Isl1 shows that Notch1^+^ radial glial fibers extend across the LGE as far as into the pial surface of ventrolateral subpallium (arrowheads) in wildtype (WT) E13.5 brain. Many Isl1^+^ cells are associated with radial glial fibers along the trajectory routes into the ventrolateral subpallium where the olfactory tubercle (OT) anlage is located (arrows, top panel). In *Nolz-1* KO brain, Isl1^+^ cells are increased in the OT-like region (arrows, bottom panel). Confocal images of the boxed regions are shown at high magnification in the far right panels. The LGE are divided into 10 equidistant bins from the dorsomedial LGE to the ventrolateral OT anlage as shown in the third panels. **G:** Isl1^+^ cells associated with Notch1^+^ radial fibers are increased in bin #9 and #10 where the mutant OT-like anlage is located compared to that in wildtype. **H:** The density of Notch1^+^ radial fibers is not changed in bin #5 to #10 of KO brains. **I:** The total number of Isl1^+^ cells associated Notch1^+^ radial fibers is not changed in KO brains. *, p < 0.05; **, p < 0.01, n =3/group. Scale bars, 200 μm in A; in C, 50 μm for 24 hr, 100 μm for 48 hr, 100 μm for 72 hr; in F, 100 μm for low magnification, 20 μm for high magnification.

Third, immunostaining of Notch1, a marker of radial glia (Gaiano et al., 2000), revealed that many Notch1^+^ radial glial fibers extended from the VZ of the LGE through the MZ into the ventrolateral subpallium where the presumptive OT anlage was located in wild-type E13.5 brains (Fig. 4G). Similar cytoarchitecture of Notch1^+^ radial glia was found in *Nolz-1* KO subpallium, and the density of Notch1^+^ radial fibers was not changed in ventral LGE of KOs (p > 0.05, n = 3, Fig. 4F, H). Because Isl1^+^ cell lineages contributed to the major part of the ventral striatum (Fig. 3F), we then investigated whether Isl1 cell lineages might migrate along radial glial fibers into the ventral striatum. Consistent with this hypothesis, in E13.5 wildtype brain, many Isl1^+^ cells were closely associated with Notch1^+^ fibers along the trajectory routes of radial glial (Fig. 4F, G), suggesting that Isl1^+^ cells generated in the LGE proper may migrate along radial glial processes into the OT anlage. For quantification, we divided the subpallium into 10 equidistant bins with the top bin #1 covering the VZ of dorsomedial LGE and the bottom bin #10 covering the ventrolateral subpallium that contained the OT anlage, and then counted the Isl1^+^ cells associated with Notch1^+^ radial glial processes in each bin. When compared to E13.5 wildtype brains, in mutants the number of Isl1^+^ cells associated with radial fibers tended to be lower from bin #1 to bin #8, but was significantly increased by 25.9% and 31.1% in bin #9 and #10, respectively, where the mutant OT analge was located (bin #9, p < 0.05, n = 3; bin #10, p < 0.01, n = 3, Fig. 4G). Because no significant changes in the total number of Isl1^+^ cells associated with Notch1^+^ fibers was found in KO LGE (p > 0.05, n = 3, Fig. 4I), these results suggested a redistribution of Isl1^+^ cells from the dorsal toward ventral LGE in *Nolz-1* KO brains.

We further attempted to clarify the identity of Isl1^+^ cells in mutant OT by double immunostaining for Isl1 and Calbindin, a marker of OT neurons at early stages (Garcia-Moreno et al., 2008). The number of Calbindin^+^ cells was not changed in E13.5 *Nolz-1* KO brains (p = 0.837, n = 3, Supplemental Fig. S8A, B). Nor was the percentage of Isl1^+^;Calbindin^+^ cells/Calbindin^+^ cells altered in KO brains (p = 0.286, n = 3, Supplemental Fig. S8A, C). Given that Isl1^+^ cells were increased in mutant OT-like regions (Fig. 4G), but Isl1^+^;Calbindin^+^ cells were not changed in mutant OT-like regions, this suggested that many Isl1^+^ cells that immigrated into the mutant OT-like regions did not differentiate into canonical Calbindin^+^ OT neurons.

Taken together, the results of these three sets of experiments, including the increases in CFDA^+^ migrating cells, pulse-labeled BrdU^E12.5^ cells and Isl1^+^ cells associated with radial glia in ventral KO striatum, consistently support the hypothesis that *Nolz-1* null mutation induces aberrant cell migration from the dorsal toward ventral striatum.

### Nolz-1 suppresses Dlx genes in the developing striatum

The formation of the abnormal ventral striatum was detected as early as E13.5, as evidenced by the expansion of the Meis2^+^ domain (Toresson et al., 2000a) in the ventral striatal anlage of *Nolz-1* KO brains (Supplemental Fig. S3A). To identify molecular mechanisms underlying the abnormal development of the mutant striatal complex, we screened a repertoire of striatum-enriched genes in *Nolz-1* KO brains. We found that Dlx homeobox genes were dramatically de-repressed in KO striatum. *In situ* hybridization showed that, in contrast to restricted expression of *Dlx1* and *Dlx2* to VZ and SVZ of wildtype striatum (Porteus et al., 1994; Price et al., 1991), *Dlx1* and *Dlx2* mRNAs were significantly up-regulated in differentiated MZ, but not in VZ/SVZ germinal zones, of E16.5 *Nolz-1* KO striatum (Dlx1, rostral, p < 0.001; middle: p < 0.05; caudal, p > 0.05, n = 3; *Dlx2*, rostral, p < 0.01; middle, p < 0.01; caudal, p < 0.01, n = 3, Fig. 5A, B). *Dlx5* was increased in MZ of rostral KO striatum (p < 0.01, n = 3, Fig. 5C). Six3, a SVZ-enriched homedomain transcription factor (Lavado and Oliver, 2011), was also markedly up-regulated in MZ of E16.5 KO striatum (rostral, p < 0.01; middle, p < 0.05; caudal, p < 0.05, n = 3, Fig. 5D). These findings suggest that Nolz-1 suppresses Dlx1, Dlx2, Dlx5 and Six3 expression in developing striatum. Consistently, we found that the majority of *Dlx1* and *Dlx2* mRNA was not co-localized with Nolz-1 protein in wildtype E13.5 striatum (Fig. 5E, F).

**Figure 5.**
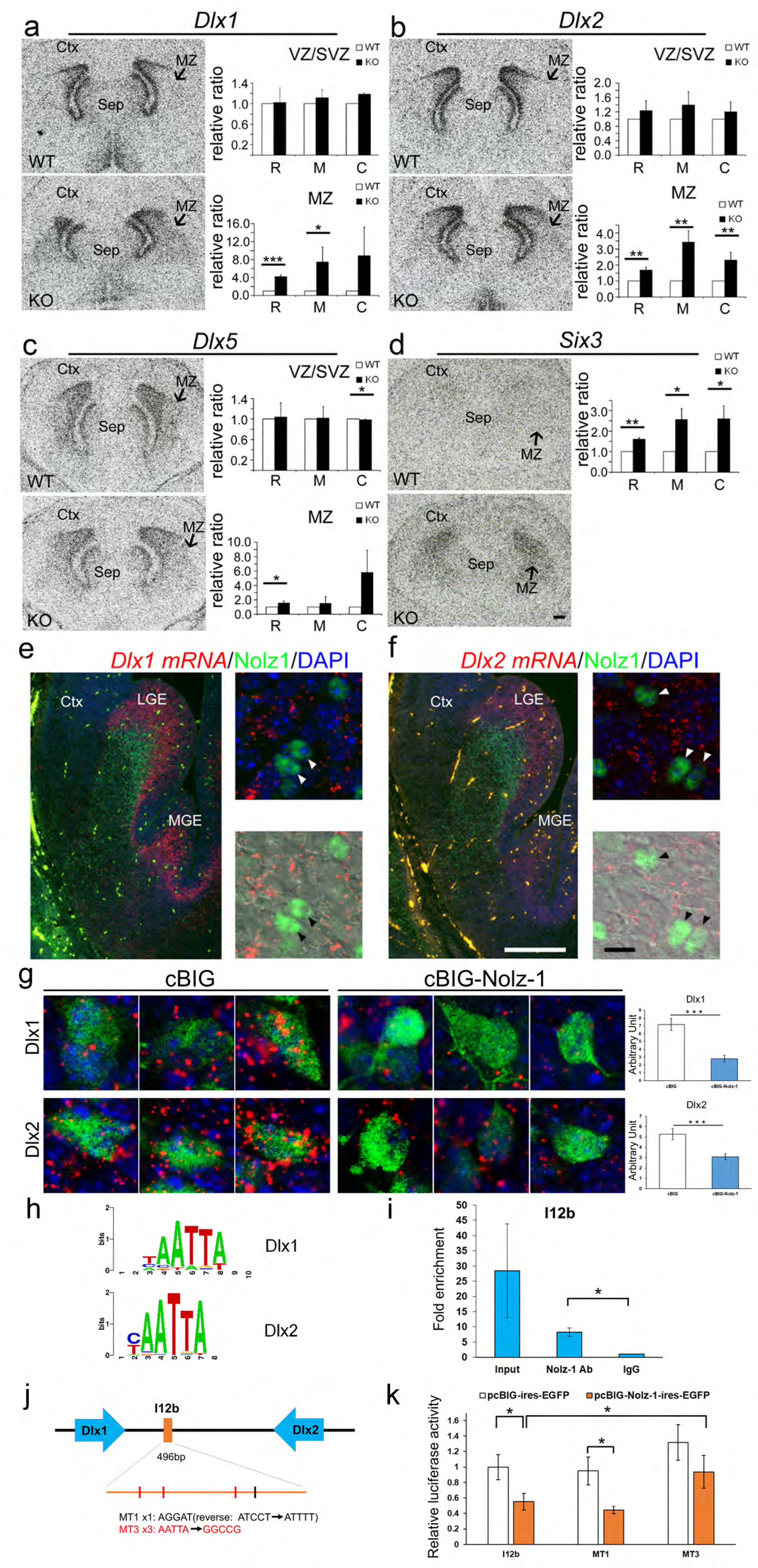
Nolz-1 directly transcriptional represses Dlx and Dlx2 in the developing striatum. **A-D:** *In situ* hybridization shows that *Dlx1* (A), *Dlx2* (B), *Dlx5* (C) and *Six3* (D) mRNAs are markedly increased in the mantle zone (MZ, arrows) of E16.5 *Nolz-1* knockout (KO) striatum compared to wildtype (WT) striatum. **E:** Double *Dlx1 in situ* hybridization and Nolz-1 immunostaining show *Dlx1* mRNA (red puncta) and Nolz-1 protein (green nuclei, arrowheads) are not co-localized in E13.5 wildtype striatum. **F:** Nor are *Dlx2* mRNA (red puncta) co-localized with Nolz-1 protein (green nuclei, arrowheads) in E13.5 wildtype striatum. **G:** Over-expression of Nolz-1 by *in utero* electroporation decreases Dlx1 and Dlx2 mRNA in the VZ and SVZ of E15.5 LGE. **H:** The ChIP-seq binding motif analysis derives a putative Nolz-1 binding motif as “AATTA” in Dlx1 and Dlx2 genes. **I:** The ChIP-qPCR assay detects a strong immunoprecipitated signal in the I12b region with Nolz-1 antibody from E15.5 striatum. **J:** Three “AATTA” motifs are located in the I12b region. **K:** The reporter gene assay shows that transfection of pcBIG-Nolz-1-ires-EGFP plasmid into E15.5 cultured striatal cells decreases pGL3-I12b-c-fos-Luc reporter gene activity. Mutations of the three motifs of “AATTA” to “GGCCG” (MT3), but not the motif of "AGGAT" to "ATTTT" (MT1), abolish the repression of reporter gene activity by Nolz-1. Ctx, cerebral cortex; LGE, lateral ganglionic eminence; MGE, medial ganglionic eminence; Sep, septum. MZ, mantle zone, SVZ, subventricular zone, VZ. ventricular zone, *, p < 0.05; **, p < 0.01; ***, p < 0.001, n =3/group. Scale bar, 200 μm in D for A-D; in F, 200 μm for low magnification, 10 μm for high magnification in E, F.

### Over-expression of Nolz-1 suppresses Dlx1/2 expression in the germinal zone of developing striatum

As *Dlx1* and Dlx2 mRNA were up-regulated in the mantle zone of Nolz-1 KO striatum (Fig. 5A, B), we tested whether over-expression of Nolz-1 could inhibit Dlx1and Dlx2 expression in the developing striatum. Indeed, over-expression of Nolz-1 by *in utero* electroporation resulted in 60.8% and 41.3% decreases in Dlx1 and Dlx2 mRNA, respectively, in the VZ and SVZ of E15.5 LGE (Fig. 5G).

### Nolz-1 represses Dlx1/2 genes through inhibition of the I12b intergenic enhancer activity

We further asked whether Nolz-1 directly transcriptionally repressed Dlx1/2 genes. It has been shown that transgenic reporter mice carrying the I12b intergenic enhancer of Dlx1/2 exhibit the reporter gene pattern similar to the endogenous Dlx1/2 pattern in telencephalon (Ghanem et al., 2003), which suggests that endogenous transcriptional regulators may act through the I12b intergenic enhancer to regulate Dlx1/2 expression. The chromatin immunoprecipitation (ChIP)-qPCR assay detected a robust immunoprecipitated signal in the I12b region with Nolz-1 antibody from E15.5 striatum (Fig. 5I), suggesting that Nolz-1 is capable of binding to the I12b intergenic enhancer *in vivo*. The ChIP-seq binding motif analysis derived a putative Nolz-1 binding motif as “AATTA” in Dlx1 and Dlx2 genes (Fig. 5H). We found three “AATTA” motifs in the I12b region (Fig. 5J). We then tested whether Nolz-1 could interact with Dlx1/2 through the I12b enhancer using a reporter gene assay. Transfection of pcBIG-Nolz-1-ires-EGFP plasmid into E15.5 striatal cell culture decreased 44.7% of pGL3-I12b-c-fos-Luc reporter gene activity compared to that of the mock control pcBIG-ires-EGFP (p < 0.05, n = 4, Fig. 5K). Moreover, mutations of the three motifs of “AATTA” to “GGCCG” (MT3) in the I12b region significantly reduced the repression of reporter gene activity by Nolz-1 (Fig. 5K). Note that a conserved zebrafish Nlz binding site “AGGAT” (Brown et al., 2009) was present in the antisense strand of the I12b region, but mutation of the "AGGAT" motif to "ATTTT" (MT1) did not affect reporter gene activity (Fig. 5K). Collectively, our findings suggest that Nolz-1 directly suppresses Dlx1/2 through inhibition of the I12b enhancer activity.

### Knocking down *Dlx1/2* gene expression alleviates the abnormal pattern of cell migration in Nolz-1 KO striatum

Dlx1 and Dlx2 genes are not only essential for tangential migration of cortical GABAergic interneurons (Anderson et al., 1997a; Marin and Rubenstein, 2003), but are also important for striatal cell migration (Anderson et al., 1997b). Because *Dlx1* and *Dlx2* were markedly up-regulated in differentiated MZ of *Nolz-1* KO striatum, we postulated that abnormal increases in *Dlx1* and *Dlx2* expression might promote aberrant cell migration. To test this hypothesis, we knocked down *Dlx1* and *Dlx2* by electroporating RNA interference (shRNAi) constructs into *Nolz-1* KO striatum to see whether this could restore normal cell migration. The knockdown effects of Dlx1 and Dlx2 shRNAi were confirmed by qRT-PCR (Supplemental Fig. S9). Dlx1 and Dlx2 shRNAi (Dlx1/2 shRNAi) and GFP reporter gene plasmids were co-electroporated into E13.5 LGE of *Nolz-1* KO brains, and brains were harvested at E18.5. The striatum was divided into 10 equidistant bins along the dorsoventral axis. Electroporation of Dlx1/2 shRNAi plasmids decreased the percentage of GFP^+^ cells in bin #7 by 34.4%, bin #8 by 46.9%, and bin #9 by 53.1% in ventral KO striatum compared to control plasmids (bin #7, p < 0.05; bin #8, p < 0.05; bin #9, p < 0.05, n = 3, Fig. 6A, C). In contrast, knockdown of *Dlx1/2* increased GFP^+^ cells by 167.2% in bin #3 in dorsal KO striatum (p < 0.05, n = 3, Fig. 6A, C). These results suggest a Dlx1/2-dependent mechanism underlying aberrant cell migration in *Nolz-1* KO striatum.

**Figure 6.**
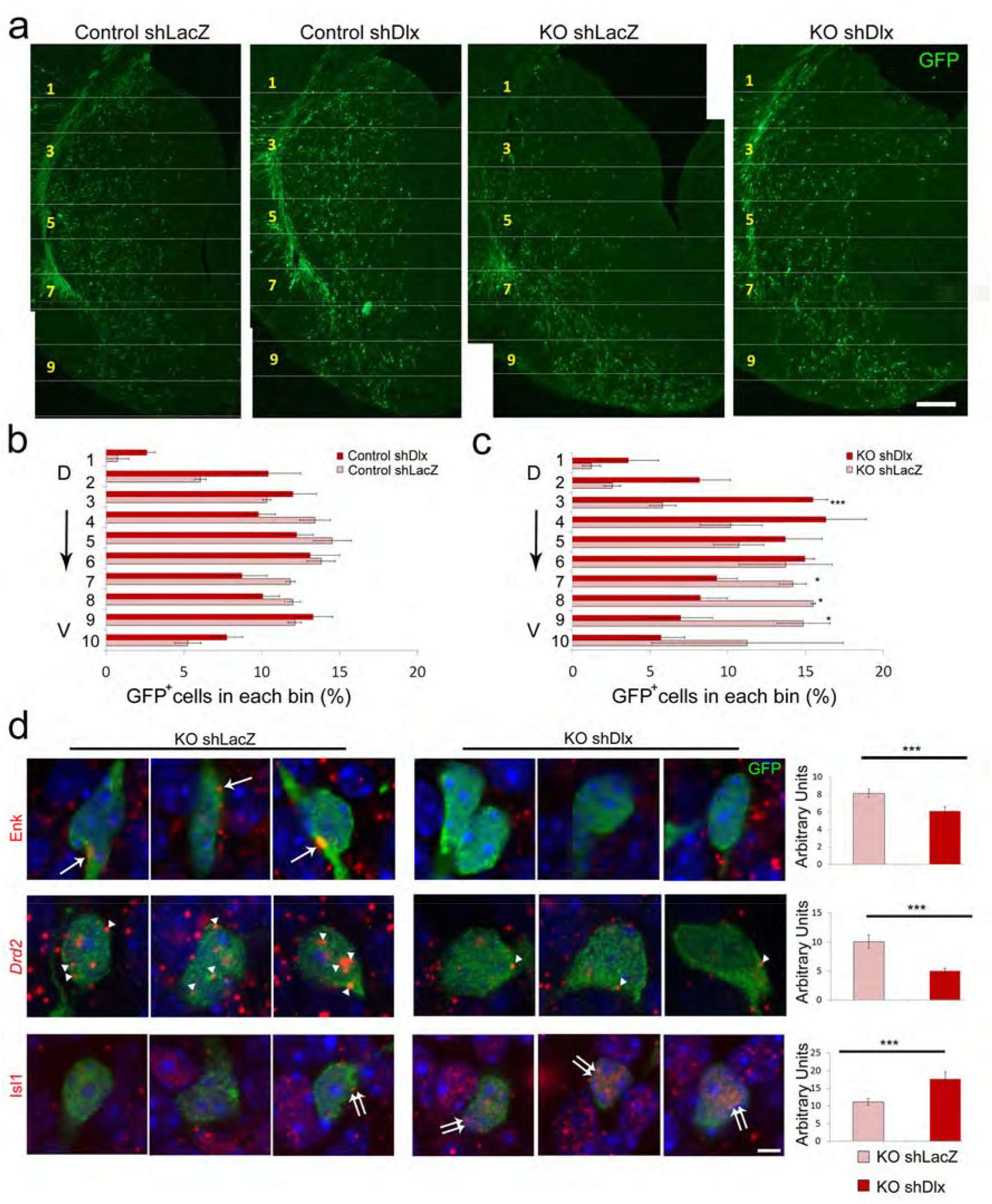
Knocking down *Dlx1/2* genes alleviates abnormal patterns of cell migration and differentiation in the *Nolz-1* knockout striatum. **A:** sh*Dlx1/2* plasmids are co-electroporated with GFP reporter plasmids into E13.5 LGE by *in utero* electroporation. The migratory pattern of sh*Dlx1/2-*GFP^+^ cells is analyzed at E18.5. In control brains, the distribution pattern of sh*Dlx1/2*-GFP^+^ cells is similar to that of mock control sh*LacZ*-GFP^+^ cells. In *Nolz-1* knockout (KO) brains, the number of sh*Dlx1/2*-GFP^+^ cells is decreased in the ventral striatum compared to that of sh*LacZ*-GFP^+^ cells. **B, C:** The striatum is divided into 10 bins along the dorsoventral axis. The percentage of sh*Dlx1/2*-GFP^+^ cells in each bin does not differ from that of sh*LacZ*-GFP^+^ cells in control brains (B). In *Nolz-1* KO brains (C), the percentage of sh*Dlx1/2*-GFP^+^ cells is decreased in the ventral striatum (bin #7-9), but is concurrently increased in the dorsal striatum (bin #3) compared to that of sh*LacZ*-GFP^+^ cells. **D:** Electroporation of sh*Dlx1/2* plasmids into E13.5 *Nolz-1* KO LGE decreases *Enk* (arrows) and *Drd2* mRNAs (arrowheads), but increases Isl1 mRNA expression (double arrows) in E18.5 KO striatum compared to the mock control of sh*LacZ*. D, dorsal; V, ventral. *, p < 0.05; ***, p < 0.001, n =3/group. Scale bars, 100 μm in A; 5 μm in C.

### Knocking down *Dlx1/2* gene expression alleviates abnormal differentiation of striatal neurons in *Nolz-1* KO striatum

Since knocking down Dlx1/2 alleviated abnormal mutant cell migration, an interesting question was whether it could also reverse abnormal increases in striatopallidal gene expression and abnormal decreases in striatonigral gene expression in *Nolz-1* KO striatum (Figs. 2, 3). Indeed, electroporation of Dlx1/2 shRNAi plasmids with a GFP reporter gene into E13.5 KO LGE resulted in a 50.4% decrease in Drd2 signals in Drd2^+^;GFP^+^ cells and a 25.1% decrease in Enk signals in Enk^+^;GFP^+^ cells in E18.5 *Nolz-1* KO striatum compared to control plasmids (Drd2, p < 0.001, n = 3; Enk, p < 0.001, n = 4, Fig. 6D). In contrast, Isl1 signals in Isl1^+^;GFP^+^ cells were increased by 59.2% in KO striatum (p < 0.001, n = 3, Fig. 6D). These results suggest that knocking down Dlx1/2 in mutant cells is able to partially reverse the aberrant differentiation of striatal neurons in *Nolz-1* KO striatum.

### Over-expression of *Dlx2* in wild-type striatum phenocopies aberrant cell migration in Nolz-1 KO striatum

If deranged Dlx1/2 signaling plays a key role in directing aberrant mutant cell migration, we reason that over-expression of Dlx1/2 in wild-type striatum may mimic the mutant migratory phenotype. We then co-electroporated a Dlx2 cDNA construct with a GFP reporter gene into E13.5 wild-type LGE. Unlike the generally even distribution of GFP^+^ cells along the dorsoventral axis of E18.5 striatum in mock controls, over-expression of Dlx2 shifted the distribution of GFP^+^ cells from the dorsal to ventral striatum. It decreased the percentage of GFP^+^ cells by 51.5% in bin # 1 of dorsal striatum (p < 0.05, n = 3, Fig. 7A, B), but increased the percentages of GFP^+^ cells by 114.7% in bin # 7 and 109% in bin #8 of ventral striatum, respectively (bin #7, p < 0.05; bin #8, p < 0.05, n = 3, Fig. 7A, B). This dorsal-to-ventral shifted pattern of cell migration was reminiscent of the aberrant pattern of cell migration found in *Nolz-1* KO striatum (Figs. 4A, B; 6A). These findings suggest that over-expression of Dlx2 is sufficient to bias cell migration from the dorsal toward ventral striatum, and further suggest that deranged Dlx1/2 signaling is likely to play a causal role in priming aberrant cell migration in *Nolz-1* KO striatum.

**Figure 7.**
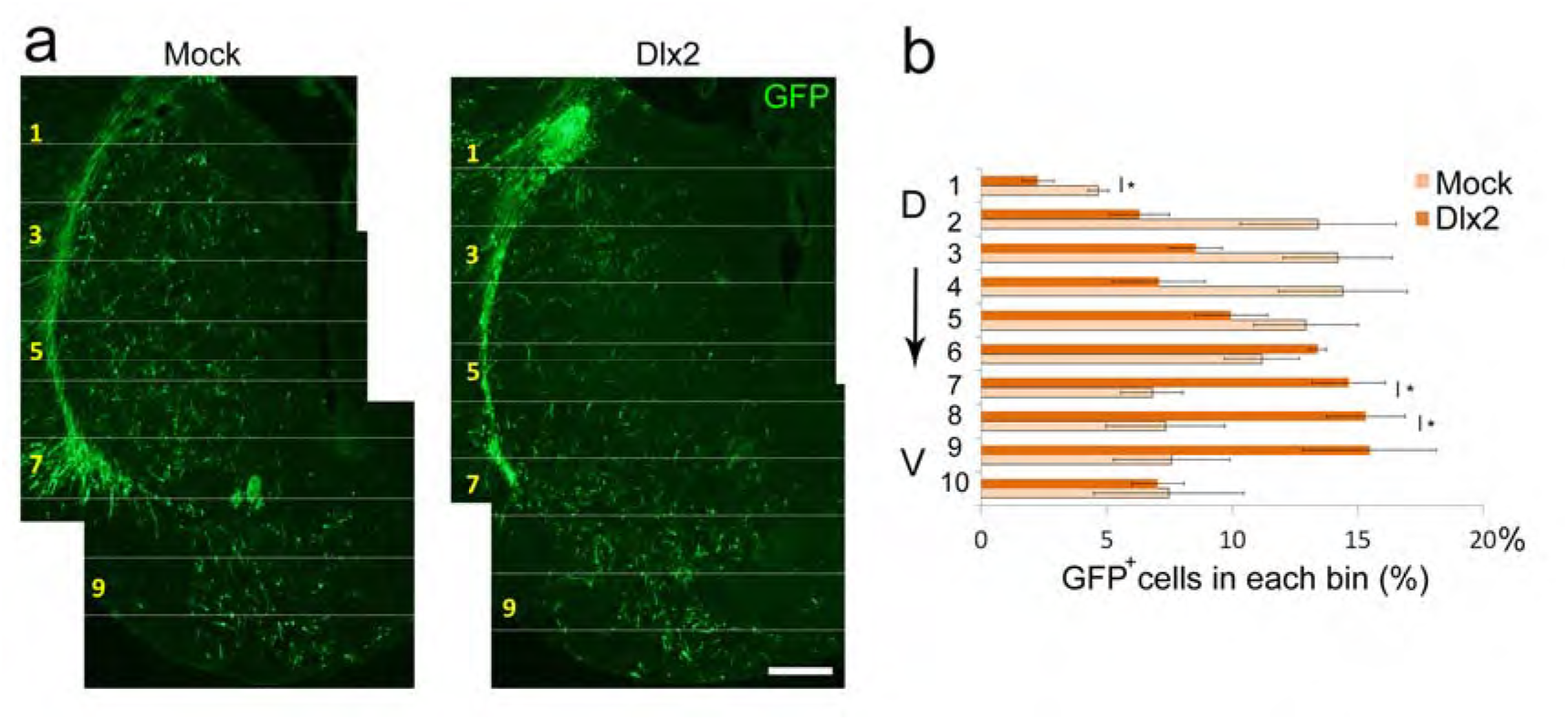
Over-expression of *Dlx2* gene in wildtype striatum phenocopies aberrant cell migration in the *Nolz-1* knockout striatum. **A:** *Dlx2* expression plasmids are co-electroporated with GFP reporter gene into E13.5 LGE by *in utero* electroporation. The migratory pattern of GFP^+^ cells is analyzed at E18.5. In mock controls, GFP^+^ cells are distributed in both the dorsal and ventral striatum. In *Dlx2* over-expressed brains, more GFP^+^ cells migrate toward the ventral striatum compared to mock control brains. **B:** The striatum is divided into 10 bins along the dorsoventral axis. Quantification shows decreased GFP^+^ cells in dorsal striatum (bin #1), but increased GFP^+^ cells in ventral striatum (bin #7-8) in *Dlx2* over-expressed brains. D, dorsal; V, ventral. *, p < 0.05, n =3/group. Scale bar, 100 μm in A. **C:** Co-transfection of Nolz-1 expression plasmid with I12b reporter gene plasmid into P19 cells suppresses luciferase reporter gene activity compared to mock control in P19 cells.

**Figure 8.**
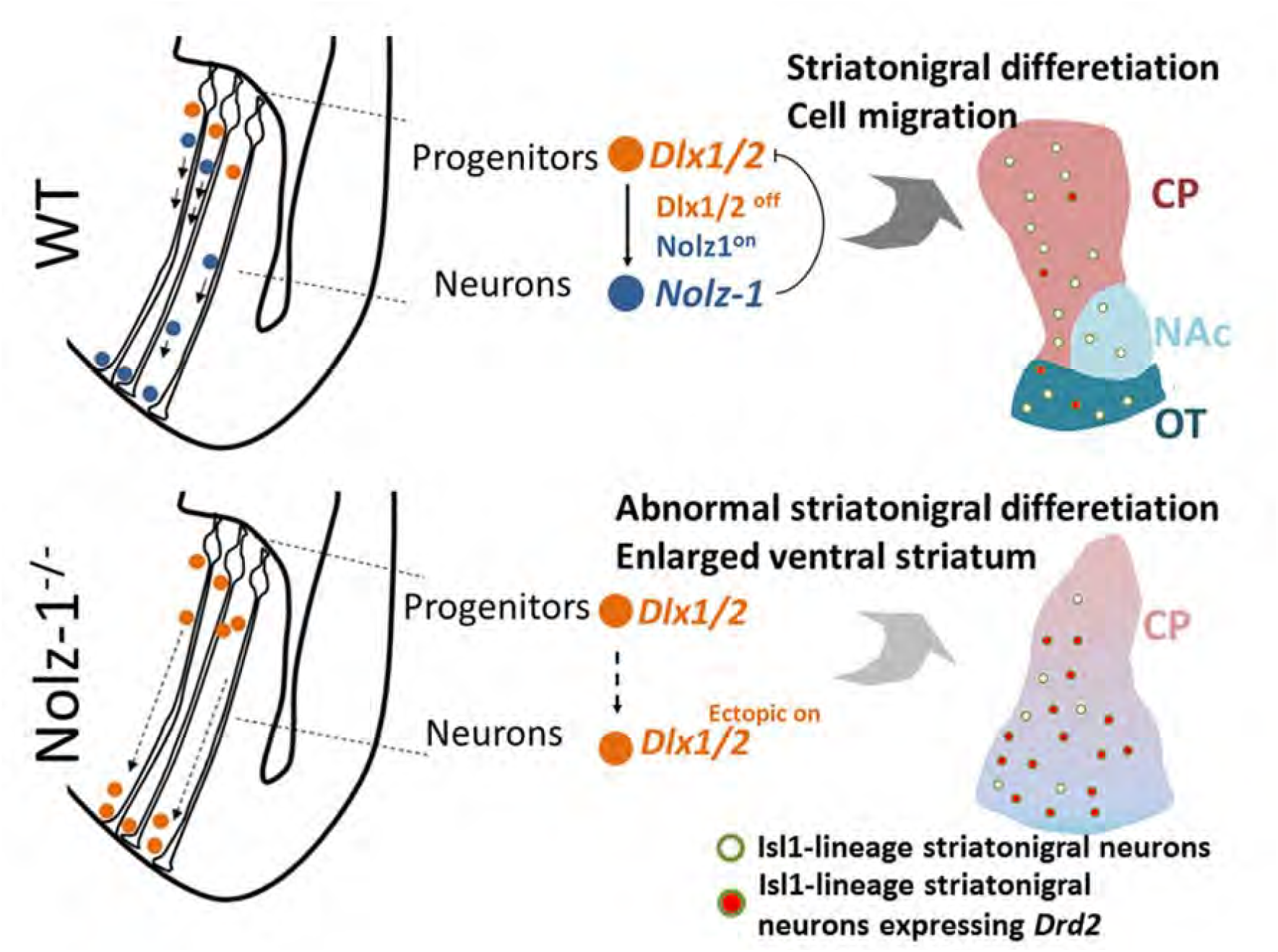
Working hypothesis. Schematic drawings showing migration and differentiation of striatal cells in striatal anlage at early (left) and late (right) stages of development. Top: In the wildtype (WT) brain, *Dlx1/2* is expressed in striatal progenitors. Repression of *Dlx1/2* by *Nolz-1* (Dlx1/2^off^;Nolz-1^on^) in postmitotic neurons is required for normal migration and differentiation of striatal neurons. Bottom: In *Nolz-1^-/-^* knockout brain, persistent expression of *Dlx1/2* (Dlx1/2^ectopic on^) in postmitotic neurons promotes aberrant cell migration from the dorsal toward ventral striatum. These postmitotic neurons are of Isl1^+^ striatonigral cell lineages, but they ectopically express striatopallidal gene *Drd2* in the absence of *Nolz-1*.

## DISCUSSION

Our present study indicates that *Nolz-1* plays an essential role for neurogenesis in the striatal complex by controlling cell migration and cell type specification. Nolz-1 directly transcriptional repress neural progenitor genes *Dlx1* and *Dlx2* to promote proper migration and differentiation of striatal neurons. N*olz-1* mutation de-represses *Dlx1/2* expression in postmitotic neurons, which results in biased neuronal migration from the dorsal to the ventral striatum, leading to altered morphogenesis of the striatal complex, and is accompanied by defects in the specification of cell type identities in CP, NAc and OT. Nolz-1, therefore, functions as a novel transcriptional regulator to control neurogenesis and allocation of striatal neurons into the dorsal and ventral striatum during development.

### Developmental origins of CP, NAc and OT

It is yet unclear whether striatal neurons in the CP, NAc and OT are derived from the same or distinct cell lineages. Much of the information on the spatiotemporal origins of CP, NAc and OT is based on analyses of mutant mice. Regarding temporal segregation of striatal progenitor cells, it has been proposed that Mash1-expressing early-born LGE cells produce some OT neurons and striosomes/patch neurons of CP, whereas Mash1 and Dlx1/2-expressing late-born cells generate other OT neurons, matrix neurons of CP and NAc neurons (Yun et al., 2003) (Kelly et al., 2018). This reasoning is based on the findings that genetic mutations of *Mash1* and *Gsh2/Gsx2* result in loss of Enk- and D2-expressing NAc and ventrolateral telencephalon, including OT in mutant brains (Casarosa et al., 1999; Corbin et al., 2000). In terms of spatial segregation of striatal progenitor cells, *Gsh2* mutant studies implicate that the dorsolateral LGE gives rise to OT neurons at early developmental stages, because the ventral striatum, particularly Isl1^+^ and Sox1^+^ OT, is reduced in *Gsh2* and *Gsh1/2* mutants in which the dorsolateral LGE cells are misspecified to pallial cells (Toresson and Campbell, 2001; Toresson et al., 2000b). Consistently, we found Notch1^+^ radial glial fibers extending from the LGE across the subpallium as far as into the OT anlage as early as E12.5, suggesting that striatal cells generated in the LGE may migrate along radial glial fibers en route through the dorsal LGE into OT at early stages of development.

### Aberrant Dlx1/2-dependent cell migration in Nolz-1 mutant brains

It has been suggested that *Gsh1/2* and *Mash1* regulate the generation of early-born striatal cells that contribute to OT, striosomes/patch compartments and NAc (Casarosa et al., 1999; Corbin et al., 2000; Toresson and Campbell, 2001; Toresson et al., 2000b; Yun et al., 2003) (Kelly et al., 2018). In line with this hypothesis, *Nolz-1* mutation resulted in the loss of the striosomal marker DARPP-32 and redistribution of BrdU^12.5^ striosomal cells from dorsal toward ventral striatum. Therefore, a hypothetical scenario is that at early stages of development, Nolz-1-dependent cell migratory mechanisms regulate early-born striatal cells that are differentially sorted into CP and OT. In the absence of *Nolz-1*, striatal cells destined for striosomes in CP are aberrantly directed toward OT.

What are Nolz-1-dependent mechanisms of cell migration? Based on the finding that knocking down aberrantly upregulated Dlx1/2 expression restored a normal pattern of cell migration in *Nolz-1* KO striatum, we propose that Nolz-1-mediated repression of Dlx1/2 in postmitotic striatal neurons plays an essential role in controlling balanced cell migration between the dorsal and ventral striatum. Dlx1 and Dlx2 are important for striatal cell migration, because double knockouts of Dlx1 and Dlx2 result in impaired migration of late-born striatal cells from SVZ into MZ, although early-born striatal cells are not affected (Anderson et al., 1997b). Dlx1 and Dlx2 are expressed by cells at the transition from proliferation to terminal differentiation in the VZ and SVZ, and they are downregulated in the MZ of striatal anlage (Porteus et al., 1994). Our present study has extended Rubenstein group’s findings by identifying Nolz-1 as a key molecule in restricting *Dlx1/2* expression to the germinal zone. Because *Nolz-1* mutation-induced upregulated expression of Dlx1/2 in the differentiated MZ causes aberrant migration and differentiation of striatal neurons, timely and precise down-regulation of Dlx1/2 in the MZ is important for proper striatal cell migration and differentiation. Rubenstein’s group has further demonstrated that Dlx1/2 promotes tangential cell migration of cortical interneurons by inhibiting axon and dendrite growth through repressing the p21-activated serine/threonine kinase PAK3 (Cobos et al., 2007). It would be of interest to see whether similar mechanisms underlie Dlx1/2-mediated migration of striatal cells.

### Cell type identity in the ventral striatum of Nolz-1 KO brains

It has been proposed that OT is a striatal-pallidal structure in which the medium-sized cell population is related to the striatum, whereas the pallidal-like cell population is invaded from the ventral pallidum (Heimer, 2003; Heimer et al., 1995). The reduction in expression of a set of ventral striatum-enriched genes, including *Acvr2a, Npy1r*, *Sox1* and *Brn4*, and the altered pattern of Nrp2 expression in ventral KO striatum (Supplemental Fig. S3B-E) do not support the possibility that dorsal striatum-derived cells differentiate into ventral striatal cell types in mutant OT-like regions. Nor do the dorsal-derived mutant cells retain the ability to differentiate into proper dorsal striatal cell types in the ventral mutant striatum, as the dorsal striatum-enriched genes, *Ebf1* and *Astn2*, remained at low levels in ventral KO striatum (Supplemental Fig. S3B). These results suggest that Nolz-1-dependent mechanisms are essential for proper specification of cell type identity in the striatal complex.

### Conditional deletion of *Nolz-1* in Isl1 cell lineages recapitulates major striatal phenotypes of *Nolz-1* KO brains

Our study shows that cell-type specific deletion of *Nolz-1* in Isl1^+^ striatonigral cell lineages using Isl1-Cre not only leads to abnormal cell migration, but also causes de-repression of the striatopallidal gene D2R and Enk in striatonigral neurons. Despite the decreases in expression of striatonigral-enriched genes and increases in expression of striatopallidal-enriched genes in striatonigral mutant neurons, it is not clear whether the mutant cells are respecified into striatopallidal neurons. Morphological changes in *Noz-1* KO striatum rendered us unable to determine whether the axons of GFP^+^ striatonigral mutant neurons were rerouted to innervate the globus pallidus. Because de-repression of Drd2 in striatonigral neurons occurs in Isl1 KO mice (Lu et al., 2014), and Isl1 is reduced in E18.5 *Nolz-1* KO striatum, *Nolz-1* mutation-induced de-repression of Drd2 in striatonigral neurons may be secondary to the loss of Isl1. Taken together, our study suggests a scenario that *Nolz-1* null mutation not only de-represses striatopallidal genes in striatonigral neurons, but also primes abnormal migration of striatonigral mutant neurons.

## MATERIALS AND METHODS

### Animals

The mice were maintained in the Animal Center of National Yang-Ming University (NYMU, Taipei, Taiwan). The protocol of animal use was approved by the Institutional Animal Care and Use Committee in National Yang-Ming University, and was conformed to the NIH Guide for the Care and Use of Laboratory Animals. For generation of *Nolz-1* knockout (KO) mice, the *Nolz-1* gene targeting vector was designed to delete the third exon of the *Nolz-1/Znf503* gene (NM_145459.3) by DNA homologous recombination. The pNolz-1-CKO-loxP-neo-TK targeting vector contained the third exon flanked with two loxP sites, a neomycin (neo)-positive selection cassette flanked with FRT site and a thymidine kinase (TK)-negative selection cassette (Supplemental Fig. S1). The generation of targeted ES cells and chimera mice was carried out by the Transgenic Mouse Core Facility at National Taiwan University. After introducing the Pvul-linearized targeting vector into 129svj ES cells and marker selection, 408 clones of ES cells were obtained. The ES cells were screened with the 3’ and 5’ probes. Three correctly targeted clones, NB2, PA5 and KA6 were obtained. These three clones of ES cells were used for blastocysts microinjection to generate chimera mice. The founder lines of chimera mice were bred with C57BL/6 mice. To generate *Nolz-1* germline knockout (KO) mice, *Nolz-1^fl/fl^* conditional KO mice of NB2 and PA5 founders were intercrossed with male protamine-Cre mice in which DNA recombinase activity of Cre was expressed in sperms. *Nolz-1^+/-^* heterozygotes were maintained by backcrossed to wild-type C57BL/6 mice. *Nolz-1^-/-^* homozygous KO mice were generated by crossing *Nolz-1^+/-^* heterozygotes. The NB2 line of *Nolz-1* KO mice was used in the present study. *Nolz-1^+/-^*, *Nolz-1^fl/+^, Isl1-Cre* (Cai et al., 2003) and *CAG-CAT-EGFP* (kindly provided by Dr. M. Colbert of Cincinnati Children’s Hospital, OH) (Burns et al., 2007) were housed in a specific pathogen-free room with 12-hr light-dark cycle in the animal center at NYMU. Transgenic mice were maintained by back-crossing with C57/BL6J strain mice. To generate *Nolz-1* conditionally knockout mice, *Isl1-Cre mice* were intercrossed with *Nolz-1^f/+^* mice. *Isl1-Cre;Nolz-1^f/+^* mice were then intercrossed with *Nolz-1^f/f^* mice. Embryonic day (E) 0.5 was defined as the day of positivity of mating plugs.

### Genotyping

The genotyping of the genetically modified mice was performed by PCR with tail genomic DNA as previously described (Lu et al., 2014). PCR genotyping primers for transgenic mice are listed as follows: Nolz-1 floxed-5’ (5′- CCAAT GTGGA AAGAT AGTAG CC-3′), Nolz-1 floxed-3′ (5′-TCCAG CAGGA AGAAG ACAGG -3′); Nolz-1 null-5’ (5′-ATTAA AGTCC CTATG TACCT AGCCC-3′), Nolz-1 null-3′ (5′- ATTTT TGGGT TCCTG GAGCC-3′). Drd2-GFP-5’ (5’-CCTAC GGCGT GCAGT GCTTC AGC-3′); Isl1-Cre-5’ (5′-ACAGC AACTA TTTGC CACCT AGCC-3′), Isl1-Cre-3’ (5′-TCCCT GAACA TGTCC ATCAG GT-3′); CAT-5’ (5′- CTGCT AACCA TGTTC ATGCC-3′), CAT-3′ (5′-GGTAC ATTGA GCAAC TGACT G-3′). The annealing temperatures are listed in the followings: 55°C for *Nolz-1* null allele, 60°C for Nolz-1 floxed allele, Isl1-Cre, and 66°C for CAG. The sizes of PCR products are follows: 550 bp for *Nolz-1* null allele, 600 bp for *Nolz-1*floxed allele, 350 bp for Isl1-Cre allele, and 300 bp for CAG allele.

### Administration of 5-bromo-2’-deoxyuridine

Timed-pregnant mice were i.p. injected with 5-bromo-2’-deoxyuridine (BrdU, 50 mg/kg) at E12.5, E13.5 or E15.5. For studying the S-phase proliferation index and the cell cycle exit index, the embryos were harvested 1 hr and 24 hr, respectively, after BrdU injection. For studying the distribution pattern of BrdU-labeled cells, embryos were harvested at E18.5. For the time course study of tracing the migration of BrdU-labeled cells, the embryos were collected 24, 48 or 72 hr after BrdU injection at E12.5.

### Preparation of brain tissue

Pregnant mice were deeply anesthetized with 3% sodium pentobarbital before abdominotomy. Embryos were removed from the uterus and kept in ice-cold PBS, and the brains were removed and immersed fixation with 4% paraformaldehyde for at least 20 hr and then cryoprotected by 30% sucrose in 0.1M PBS for 2 days. Brains were cut at 12-18 μm by a cryostat (Leica).

### Dissection of embryonic brains

E13.5 brains of C57BL/6 mice were dissected out from the head, and were then embedded in 4% low-melting agarose for sectioning (Invitrogen). Fresh brain slices were sectioned at 300 μm by vibratome. Brain slices with the proper anatomical level were used for further dissection. The pallidum appeared darker under the dissecting microscope, and it was dissected away from brain slices. The septum was cut off as well. The remaining striatal complex anlage was cut in half.

The dorsal and ventral parts of the striatum were collected and frozen with dry ice for microarray and qRT-PCR analyses..

### Genome-wide microarray analysis

Total RNA was extracted from the dorsal or ventral parts of E13.5 LGE by Trizol^®^ according to the manufacture’s instruction. The genome-wide microarray analysis was performed with Affymetrix mouse 430A gene chips by custom service of NYMU Genomic Center. Genes with the expression levels higher or lower than 1.3 fold between the dorsal or ventral striatum were selected for functional categorization by the IPA analysis (Qiagen).

### Anterograde axonal tracing

After fixation in 4% paraformaldehyde, E18.5 brains were sectioned using a vibratome at a thickness of 100 μm. Small crystals of DiI (Invitrogen) were placed in the anterior part of the striatum with a fine glass pipette. The DiI-labeled brain slices were stored in 4% paraformaldehyde in the dark for ~2 months at 37°C. The DiI-labeled brains were sectioned at a thickness of 100 μm and counterstained with DAPI.

### Tracing cell migration with CFDA in living embryonic brains

The cell tracker CFDA (5 mM in DMSO, Invitrogen) was mixed with the fast green dye (1% in PBS; 1:9, v/v). Microinjections of ~100-200 nl of the CFDA solution aiming to target the dorsal part of LGE was performed in living E14.5 embryos with the aid of an ultrasound image-guided microinjection system (S-Sharp Corporation, New Taipei City, Taiwan) and with empirical experience. The brains were harvested ~31 hr after surgery. We injected 45 control and 12 *Nolz-1* knockout embryos. The location of CFDA injection site in the LGE was identified by histology. The successful CFDA injection site was defined as ~400 μm-radius circle centering at the intensive CFDA-labeled ventricular zone of dorsal LGE. Successful CFDA injections with the injection sites specifically restricted to the dorsal part of LGE were analyzed in 3 controls and 3 knockout brains.

### Electroporation plasmids

The pUS2-GFP and pUS2-Dlx2 plasmids were kindly provided by Dr. J-Y Yu (NYMU, Taiwan). The pKLO.1-TRC shRNA constructs for shLacZ (TRC0000072231), shDlx1 (TRC0000070588) and shDlx2 (TRC0000321374) were obtained from the RNAi Core Facility (Academia Sinica, Taiwan).

### shDlx1 and shDlx2 knockdown efficiency test

P19 cells were cultured as previously described (Chang et al., 2004). The shDlx1, shDlx2 or shLacZ plasmids were transfected into P19 cells with Lipofectamine LTX reagent (Invitrogen). 24 h after transfection, the P19 cells were selected with 10 mg/ml puromycin (Sigma) for 48 h with a change of fresh medium containing puromycin every day. The knockdown efficiency was measured by quantitative RT-PCR using the following primers: Dlx1-5’ (5’-GTGCT TGATT ACAGA GGTCT CCCTG-3’), Dlx1-3’ (5’-TTGCC CGACA CGGGG CTGTT GAGAC-3’); Dlx2-5’ (5’-GGCTG AGAAC TAAAT CCAGG CA-3’), Dlx2-3’ (5’-GGGAC AGGAA AGAGC ACGGG TG-3’).

### *In utero* electroporation

The pUS2, pUS2-Dlx2, pUS2-GFP and shRNAi plasmids were prepared using the endotoxin-free plasmid preparation kits (Qiagen, NovelGene). For the experiment of *Dlx2* over-expression, the pUS2-GFP plasmid was mixed with pUS2 control or pUS2-Dlx2 plasmids (0.2:2, w/w). For the *Dlx1/2* knock-down experiment, the pUS2-GFP plasmid was mixed with shLacZ or shDlx1/2 plasmids (0.2:3, w/w). For the experiment of Nolz-1 over-expression, pcBIG-Nolz-1-ires-GFP and pcBIG-ires-GFP plasmids were used. 1.5 μl of the plasmids were injected into the lateral ventricle of E13.5 forebrains of mouse embryos followed by electroporation (two trains of 4 pulses, 40V, 30 ms, interval: 970ms BTX ECM 830). The electroporated brains were harvested at E18.5 for histological analysis except that Nolz-1 over-expressed brains were harvested at E15.5.

### Immunohistochemistry

Antigen retrieval for immunohistochemistry was performed for Isl1 immunostaining [(3A4, 4D5 antibodies) with 10 mM citric acid buffer (pH 6.0) at 95°C, 15 min] and BrdU Immunostaining [1 N HCl/0.1M PB at 45°C, 30 min, 0.1 M borated buffer (pH 8.6), 10 min]. Sections were washed with 0.1 M PBS twice for 5 min, 0.1% Triton X-100 in 0.1M PBS for 5 min and 3% H_2_O_2_/10% methanol in 0.1 M PBS for 5 min. Sections were incubated with the following primary antibodies overnight at room temperature: monoclonal rat anti-BrdU antibody (1:500, Accurate), polyclonal rabbit anti-Brn4 (1:500; kindly provided by Dr. E.B. Crenshaw III of University of Pennsylvania), polyclonal rabbit anti-calbindin antibody (1:1,000, Swant), polyclonal rabbit anti-CalDAG-GEFI (1:2000; kindly provided by Dr. A.M. Graybiel of MIT), polyclonal goat anti-D1R antibody (1:2,000, Frontier Institute, Japan), polyclonal goat anti-D2R antibody (1:2,000, Frontier Institute, Japan), polyclonal rabbit anti-DARPP32 (1:2000; Cell Signaling), monoclonal mouse anti-Foxp1 antibody (1:1,000; kindly provided by Dr. J. Cordell of John Radcliffe Hospital, Oxford, UK), polyclonal rabbit anti-Foxp2 antibody (1:1,000; Abcam), polyclonal chicken anti-Green Fluorescent Protein (GFP; 1:1000; Abcam), polyclonal rabbit anti-Isl1 antibody (1:1,000, Abcam), monoclonal 3A4 or 4D5 mouse anti-Isl1 antibody (1:100, Developmental Studies Hybridoma Bank), monoclonal mouse anti-Ki67 (1:500 BD Pharmingen^TM^), polyclonal rabbit anti-met-enkephalin antibody (1:5,000, ImmunoStar), polyclonal rabbit anti-Nolz-1 antibody (1:1,000) (Ko et al., 2013), monoclonal rabbit anti-Notch1 antibody (1:1,000, Abcam), polyclonal rabbit anti-Neuropilin-2 antibody (1:2,000, Cell Signaling), polyclonal goat anti-Plexin D1 antibody (1:1,000, R&D system), polyclonal rabbit anti-phospho-Histone H3 antibody (1:1,000, Millipore), monoclonal rabbit anti-Sox1 antibody (1:200; Epitomics), polyclonal rabbit anti-substance P antibody (1:3,000, Eugene Tech Inc.). After incubation with primary antibodies, the sections were incubated with appropriate secondary antibodies (1:500) for 1 hr: biotinylated goat anti-rabbit antibody, horse anti-mouse antibody, rabbit anti-goat antibody and rabbit anti-rat antibody. The sections were then processed with avidin-biotin based immunostaining (Vectastain Elite ABC kit, Vector Laboratories). Avidin-FITC (Vector Laboratories) or the tyramide Amplification system (TSA, PerkinElmer) were used to detect signals. Immunostained sections were counterstained with DAPI (Molecular Probes).

### *In Situ* hybridization

The brain sections of wild-type and littermate mutant mice were mounted on the same slides to ensure they were processed under the same condition. The mouse *Dlx1* riboprobe (126 bp, gift of Dr. D. Duboule of the European Molecular Biology Laboratory, Heidelberg, Germany), mouse Dlx2 riboprobe (363-1361 bp, gift of Dr. J.-Y. Yu of National Yang-Ming University, Taiwan), rat *Drd2L* riboprobe (128-1372 bp, gift of Dr. K. Kobayashi of Fukushima Medical University, Fukushima, Japan) were used. The rat *Dlx1* (S81923, 1-132 bp), rat *Dlx2* (S81930, 17-236 bp), rat *Dlx5* (S81929, 30-296 bp), mouse *Mash1* (NM_001002927, 327 bp), *Astn2* (NM_019514.3, 671-1478 bp), *Acvr2a* (NM_007396.4, 1752-2722 bp), *Ebf1* (NM_053820, 1847-2214 bp), *Npy1r* (NM_010934.4, 566-1455 bp), *Penk* (NM_001002927.2, 312-1128 bp) and mouse *Six3a* (D83144.1, 1495-1773 bp) were cloned by PCR and then subcloned into pCRII or pGEM-T-easy vectors (Promega). *In situ* hybridization was performed with ^35^S-UTP-labeled or DIG-labeled probes as previously described (Chang et al., 2004; Liao et al., 2005; Lu et al., 2014).

### TUNEL assay for cell death

TUNEL staining was performed using the *In Situ* Cell Death Detection Kit (Roche) according to the manufacture’s instruction. Sections were rinsed 5 minutes twice with 0.1 M PBS, and then incubated in 0.1 M glycine/PBS for 30 min at room temperature. After dipping with 0.1 M PBS briefly, sections were incubated in the blocking solution containing 3 % H_2_O_2_ in methanol for 10 min at room temperature and were then treated with the permeabilization solution (0.1 % sodium citrate, 0.1 % Triton-X100 in H_2_O) for 2 min on ice. After three times with 0.1 M PBS for 5 minutes, sections were incubated in the label solution containing 1,500 unit/ml terminal transferase (TdT) for 1 hr at 37 °C to label free 3’OH ends in fragmented DNA with fluorescein-dUTP. Sections were counterstained with DAPI.

### Quantitative RT-PCR

Striatal tissue was dissected from E18.5 brains. Total RNA extraction and quantitative RT-PCR for *Drd1a, Isl1, Ebf1, substance P, PlexinD1, Chrm4, Slc35d3, Drd2, preproenkephalin, A2aR, Gpr6* and *GAPDH* were performed as previously described (Lu et al., 2014). The real-time PCR reactions using Syber green dye-based detection were performed by Applied Biosystems StepOnePlus System. Acvr2a-5’ (5’-CTCGT TGCAC TGCTG CAGAT-3’), Acvr2a-3’ (5’-AGAGA TGGAT GCTGG CCAAT-3’); Astn2a-5’ (5’-CTCTG TATGC TGTGG ATACT CG-3’), Astn2a-3’ (5’-TCCTT GCCAC TTGTG TAACC-3’); Calb1-5’ (5’-CAGAA TCCCA CCTGC AGTCA-3’), Calb1-3’ (5’-TCCGG TGATA GCTCC AATCC-3’); Ebf1-5’ (5’-GGCCT CTGAG AGTGG AGCAA-3’), Ebf1-3’ (5’-GAGTG CGAGC GAGAT TTTGG-3’); Fgf14-5’ (5’-GAAGT TGCCA TGTAC CGAGA AC-3’), Fgf14-3’ (5’-ATGTG GTCTT GCACT TGTTG AC-3’); Fgf15-5’ (5’-GGAGG ACCAA AACGA ACGAA-3’), Fgf15-3’ (5’-TGACG TCCTT GATGG CAATC-3’); Gli1-5’ (5’-ATGCT CCGTG CCAGA TATGC-3’), Gli1-3’ (5’-GGGTA CTCGG TTCGG CTTCT-3’); Nolz-1-5’ (5’-TTGAA GCTGA GCGAC ATC-3’), Nolz-1-3’ (5’-CGGC TCCTT CTTAT CTGAA-3’); Npy1r-5’ (5’-CCTGC CCTTG GCTGT GATAT-3’), Npy1r-3’ (5’-AAGAA TAATC ACCGC CCCAT AA-3’); Nrp2-5’ (5’-GTGGA CCTGC GCTTC CTAAC-3’), Nrp2-3’ (5’-TGCCG GTAGA CCATC CAATC-3’); Nts-5’ (5’-GAGCC CTGGA GGCAG ATCTA-3’), Nts-3’ (5’-GACGG AGGAC TTGCT TTGCT-3’); Otx2-5’ (5’-AAGCA GAGGT GATCC GGTGT T-3’), Otx2-3’ (5’-CGTGA ATGAG CCTGG GACAT-3’); Plxnc1-5’ (5’-GTGAT CCGGG TTAGC CATGT-3’), Plxnc1-3’ (5’-GCGGC TTGAT GGAAG TGAGA-3’); Wnt-7b-5’ (5’-GCAGT GTGGA TGGAT GTT-3’), Wnt-7b-3’ (5’-GGCTA CCCAG TCGTT AGTA-3’); Zic1-5’ (5’-CTGGC GTTCA CATTC CTCTA TG-3’) and Zic1-3’ (5’-ACATC TCAGC CCCCC AATAA A-3’).

### Chromatin immunoprecipitation (ChIP) assay

The LGE was dissected form E15.5 ICR mice. The tissue was processed using the EZ-Magna ChIPTM A/G kit (EMD Millipore) according to the manufacturer’s instructions. The reagents and spin columns were used in the following steps that were provided from the kit. In brief, 1/10 volume of fresh 11% formaldehyde solution was added to culture plates for 15 min at room temperature followed by quenching formaldehyde with 1/20 volume of 2.5 M glycine. After washing twice with 5 ml PBS, the cells were harvested by spinning at 1,350 x RPM for 5 min at 4°C. The cells were re-suspended in 0.5 ml cell lysis buffer containing 2.5 μl of protease inhibitor cocktail II. The cells was spun down at 800 g for 5 min at 4°C. The pellet was re-suspended in 0.5 ml nuclei lysis buffer and subject to sonication to shear DNA with a sonicator (30s on, 30s off, 15 cycles, Bioruptor UCD-200). The debris was removed by centrifuge at 13,000 rpm for 10 min at 4°C. Rabbit anti-Nolz-1 FLP2 antibody (5 μg) (Ko et al., 2013) or rabbit IgG (5 μg) were preabsorbed with primary neuron for 1 hr then captured by protein A/G Magna beads (20 μl). The antibody-bead complex and the sheared DNA lysate were incubated overnight at 4°C. The antibody-DNA complexes were washed sequentially with low salt, high salt, LiCl and TE buffers followed by incubating in the ChIP elution solution containing proteinase K for 2 hr at 65°C. DNA purification was performed using purification spin columns. The purified DNA products were used as template for qPCR reaction with I12b primers: (5′-GCTGCCATCTCTGCAATTCTC-3′) and reverse (5′-GCTCCATACACCTGCAAGCA-3′), which was carried out with the Applied Biosystems StepOne system. The PCR reaction solution containing template DNA, FMR157 primers and 2X TaqMan^®^ Fast Universal PCR Master Mix (Applied Biosystems®). The PCR reactions were performed with the following conditions: 95°C for 10 min followed by 40 cycle at 95°C for 10 sec, 60°C for 1 min.

### ChIP sequencing (ChIP-seq)

The chromatin immunoprecipitation (ChIP) was performed on E15.5 striatal tissue with anti-Nolz-1 FLP2 antibody. The ChIP sequencing (ChIP-seq) was carried out by NYMU Genome Research Center. The ChIP-seq data were analyzed by the PscanChIP web application to search for over represented sequence motifs according to the motif descriptors of the JASPAR databases.

### Reporter gene assay

The genomic region of mouse I12b enhancer (496 bp) was amplified by PCR with the primers (sense: ATGCTAGCCGCGTACAGCTGCAAACCCA, antisense: ATGCTAGCACCAGTGAGGGAAAGGTTGG). The amplicons of I12b were constructed into the pGL3-cfos-Luc plasmid by NheI site. There is no conserved Nlz binding motif "AGGAT" (Brown et al., 2009) except that a reverse sequence "ATCCT" is located in the antisense strand in I12b region. Based on our binding motif analysis with ChIP-seq, a putative Nolz-1 binding motif was "AATTA", which was present in the I12b region. The I12b DNA that contained the mutation sites ["ATCCT" to "ATTTT" (MT1) or “AATTA” to “GGCCG” (MT3)] was obtained by nucleic acid synthesis (MISSION BIOTECH company, Taiwan). Striatal tissue was dissected from the brains of E15.5 embryos of ICR mice. Tissues were washed three times with PBS containing glucose and then digested with the dissociation solution containing papain (10.2 units/ml, Sigma), 0.5 mM EDTA, 1.5 mM CaCl_2_ and cysteine (0.2 mg/ml) in PBS for 30 min at 37 °C, followed by adding 0.02% DNAase I (Sigma) and 1.1 mM MgCl_2_ into the dissociation solution and incubated for additional 5 min. Enzymatic digestion was stopped by adding 2% Fetal bovine serum (FBS, Thermo Fisher Scientific). The digested tissue was collected by centrifugation and mechanically dissociated in Neurobasal medium (Thermo Fisher Scientific) supplemented with 1X B27 (Thermo Fisher Scientific). The cells were plated at the density of 10^6^ cells per well in 24-well culture plate pre-coated with poly-D-lysine. Striatal cell culture was maintained with 5% CO_2_ at 37°C. The pGL3-I12b-c-fos-Luc, pGL3-MT1-c-fos-Luc or pGL3-MT3-c-fos-Luc plasmids were co-transfected with pcBIG-Nolz-1-ires-GFP or pcBIG-ires-GFP plasmids and pGL4-Renilla plasmids [plasmid ratio 1:1:0.1 (w:w:w), 0.5 ug/24 well per well] into cultured striatal cells using Lipofectamine LTX with Plus Reagent (Thermo Fisher Scientific). Five days after transfection, cells were lysed to measure luciferase activity using the Dual-Luciferase Reporter Assay System (Promega). The luminescence signal was detected with a luminescence Reader (VICTOR^TM^ x2, PerkinElmer).

### Image analyses, quantification and statistics

Photomicrographs of immunostained brain sections were taken under fluorescence microscopes (Olympus BX51, BX63) or confocal microscope (Zeiss LSM700). Brightness adjustments for the whole field of photomicrographs were made, and final plates were composed using Adobe Photoshop CS (Adobe Systems). Quantification of cell number and area was performed with the aid of Image J software. For the M-phase index, the number of PH3^+^ cells was normalized with the length of VZ or the area of SVZ germinal zone. For quantification of the area of the striatal complex, Foxp1 immunostaining was used to define the striatal complex. A line from the bottom of the lateral ventricle to the piriform cortex was drawn to separate the dorsal from the ventral parts of the striatum. For quantification of CFDA-labeled migrating cells, the percentage of CFDA^+^ cells in each zone of the concentric rings (200 μm apart) was calculated by dividing the number of CFDA^+^ cells in each zone with the total number of CFDA^+^ cells. For quantification of Notch1^+^ radial glial fibers and Isl1^+^ cells associated with radial glial fibers, we divided the subpallium into 10 equidistant bins along the plane of radial glial fibers with the top bin #1 covering the ventricular zone of dorsomedial LGE and the bottom bin #10 covering the ventrolateral subpallium containing the OT anlage. The numbers of radial glial fibers and Isl1^+^ cells associated with radial glial fibers in each bin were counted. For calculating the percentage of BrdU^E12.5^ cells 24, 48, and 72hr after BrdU injection, the number of BrdU^E12.5^ cells in each equidistant zone of the concentric rings (100 μm apart) were divided by the total number of BrdU^E12.5^ cells in the striatum. For quantification of the distribution pattern of electroporated GFP^+^ cells in the striatum, the striatal complex was divided into ten equidistant bins along the dorsoventral axis. The number of GFP^+^ cells in each bin was divided by the total number of GFP^+^ cells to derive the percentage of GFP^+^ cells in each bin. For quantitative analysis of *in situ* hybridization with ^35^S-UTP-labeled probes, photomicrographs of autoradiographic X-ray films were taken. The 35S-UTP-labeled signal intensity in the autoradiograms was analyzed using the Image J software. For quantification of double *in situ* hybridization with Dig-labeled probes and immunostaining, photomicrographs of confocal images were taken (Zeiss LSM700). Fluorescent Dig-labeled signal intensity in GFP^+^ cells was measured using the Image J software. *Student’s t*-test was used for statistical analysis.

## ACKNOWLEDGMENTS

We thank Drs. D. Anderson, M. Colbert, J. Cordell, E.B. Crenshaw III, D. Duboule, A.M. Graybiel, K. Kobayashi and J.-Y. Yu for providing the reagents and transgenic mice, and the Transgenic Mouse Model Core Facility of NRPB at NTU for help in the generation of *Nolz-1* mutant mice. This work was supported by National Science Council grant NSC95-3112-B-010-014, NSC96-3112-B-010-007, NSC97-3112-B-010-005, NSC99-2311-B-010-005-MY3, NSC101-2321-B-010-021, NSC102-2321-B-010-018, Ministry of Science and Technology grant MOST103-2321-B-010-009, MOST104-2311-B-010-010-MY3, MOST-107-2321-B-010-002, and MOST107-2320-B-010-041-MY3 (F.-C.L.).

## AUTHOR CONTRIBUTIONS

K.-M.L., S.-Y.C., H.-A.K., T-.H.H., J.H.-J.H., Y.-T.Y., S.L.-Y.C., F.-C.L. designed research; K.-M.L., S.-Y.C., H.-A.K., T-.H.H., J.H.-J.H., H.-Y.Y, S.L.-Y.C. performed research; S.E. contributed transgenic mice, edited manuscript; K.-M.L., S.-Y.C., H.-A.K., T-.H.H., J.H.-J.H., Y.-T.Y., S.L.-Y.C., F.-C.L. analyzed data; F.-C.L wrote the paper.

## SUPPLEMENTARY FIGURES

**Supplementary Figure 1.**
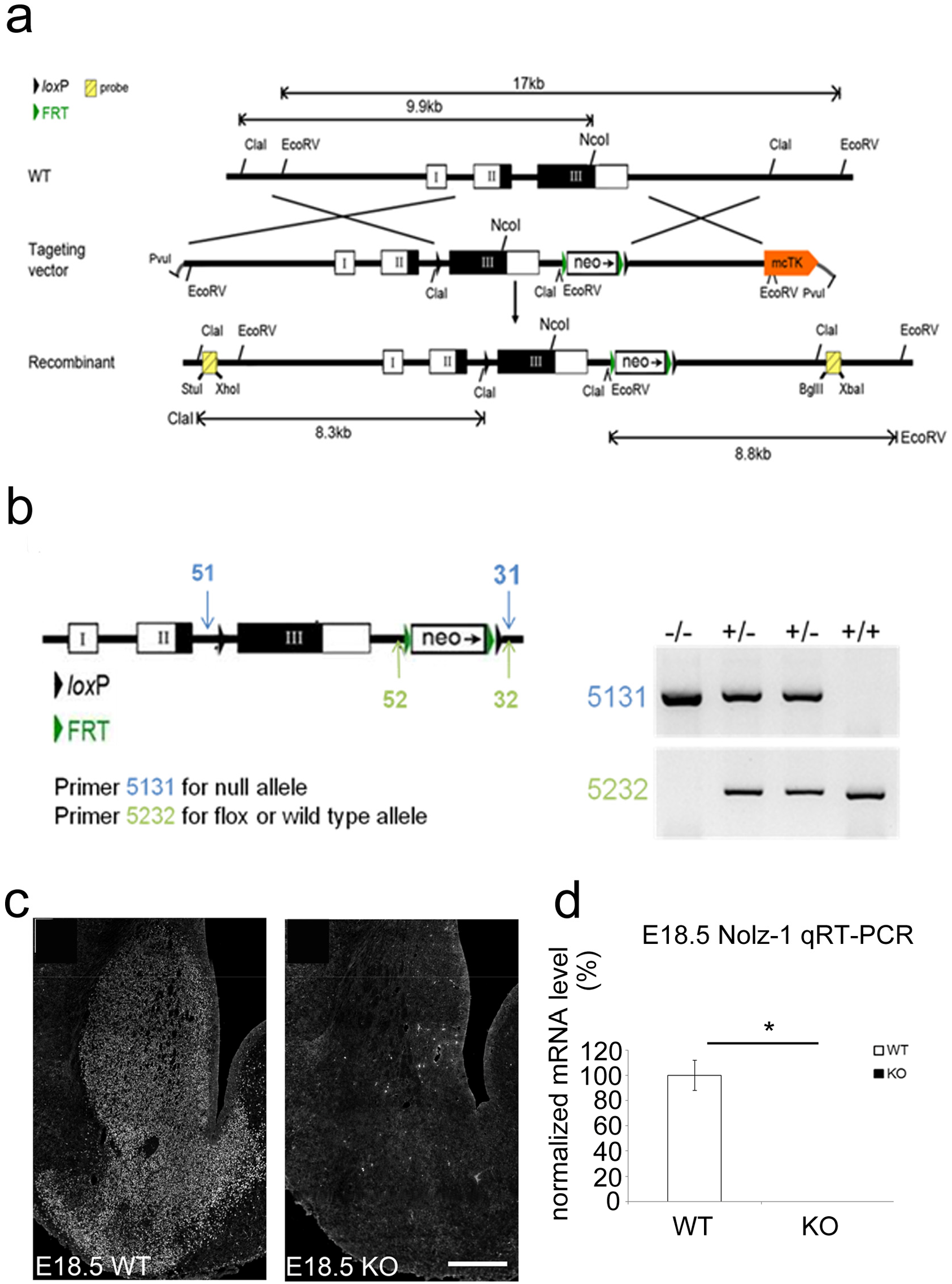
Generation of *Nolz-1* knockout mice. **(a)** The design of Nolz-1 targeting vector and the strategy of homologous recombination. **(b)** The PCR primers sites for genotyping *Nolz-1* wildtype and null alleles. **(c, d)** Deletion of *Nolz-1* gene is confirmed at the protein level by immunostaining **(c)** and at the transcript level by qRT-PCR **(d)** in E18.5 Nolz-1 knockout striatum. *, p < 0.05. n = 3/group. Scale bar, 200 μm.

**Supplementary Figure 2.**
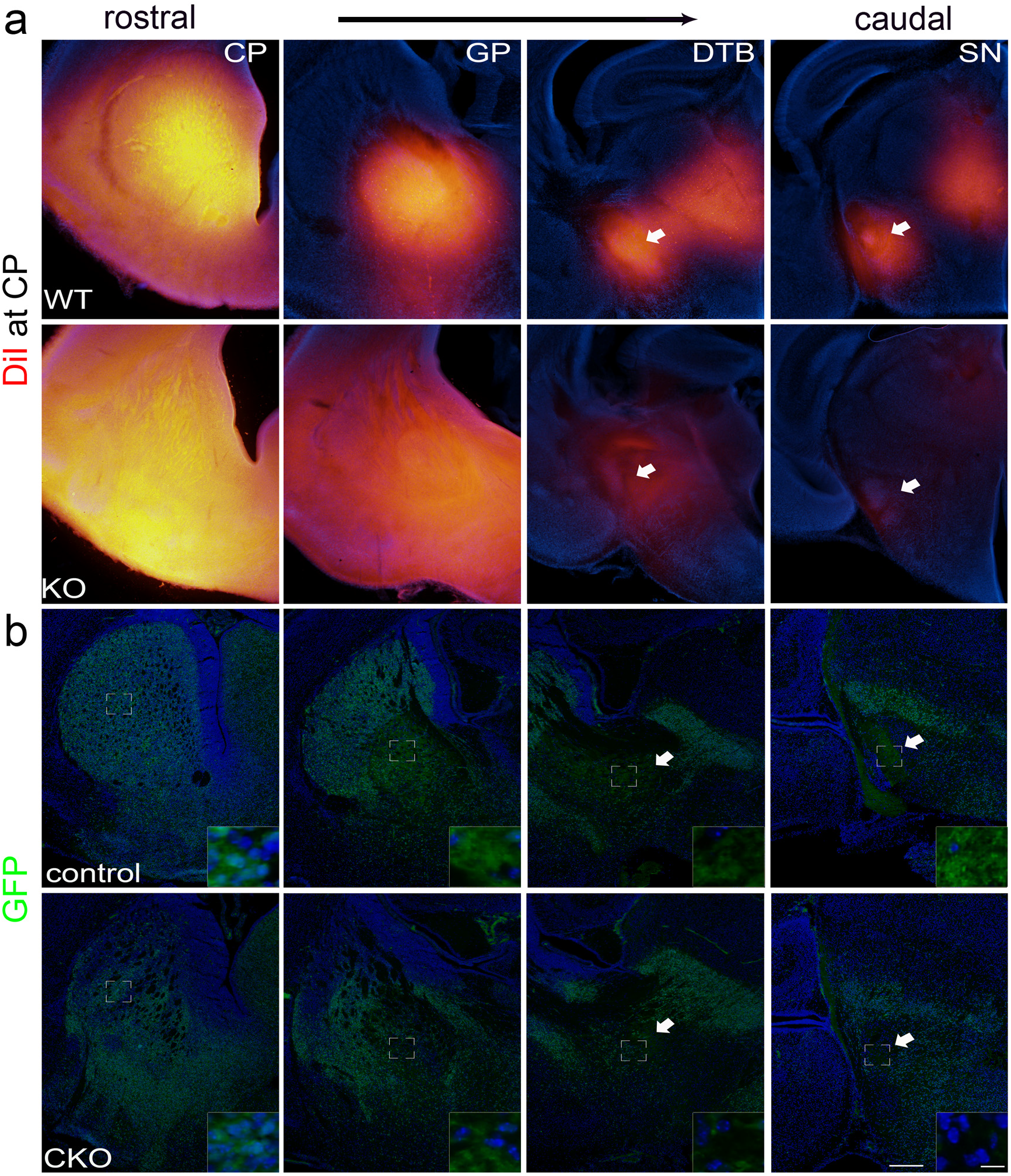
Defective striatofugal axonal projections in Nolz-1 knockout brains. **(a)** Anterograde DiI tracing shows that DiI-labeled axons extend from the caudoputamen (CP) along the globus pallidus (GP), cerebral peduncles at diencephalon and telencephalon boundary (DTB) into the substantia nigra (SN) (arrows, top panel) of wildtype (WT) brain. DiI-labeled striatal axons were markedly decreased in the cerebral peduncles at DTB and SN of Nolz-1 knockout KO) brain (arrows, bottom panel). **(b)** Similarly defective striatonigral pathway is also observed in the Isl1-Cre;Nolz1^fl/fl^;CAG-CAT-EGFP conditional KO (CKO) brains in which significant reduction of striatonigral GFP^+^ axons is found along the routes of striatonigral axonal projections. Scale bars, 200 μm for low magnification, 10 μm for high magnification.

**Supplementary Figure 3.**
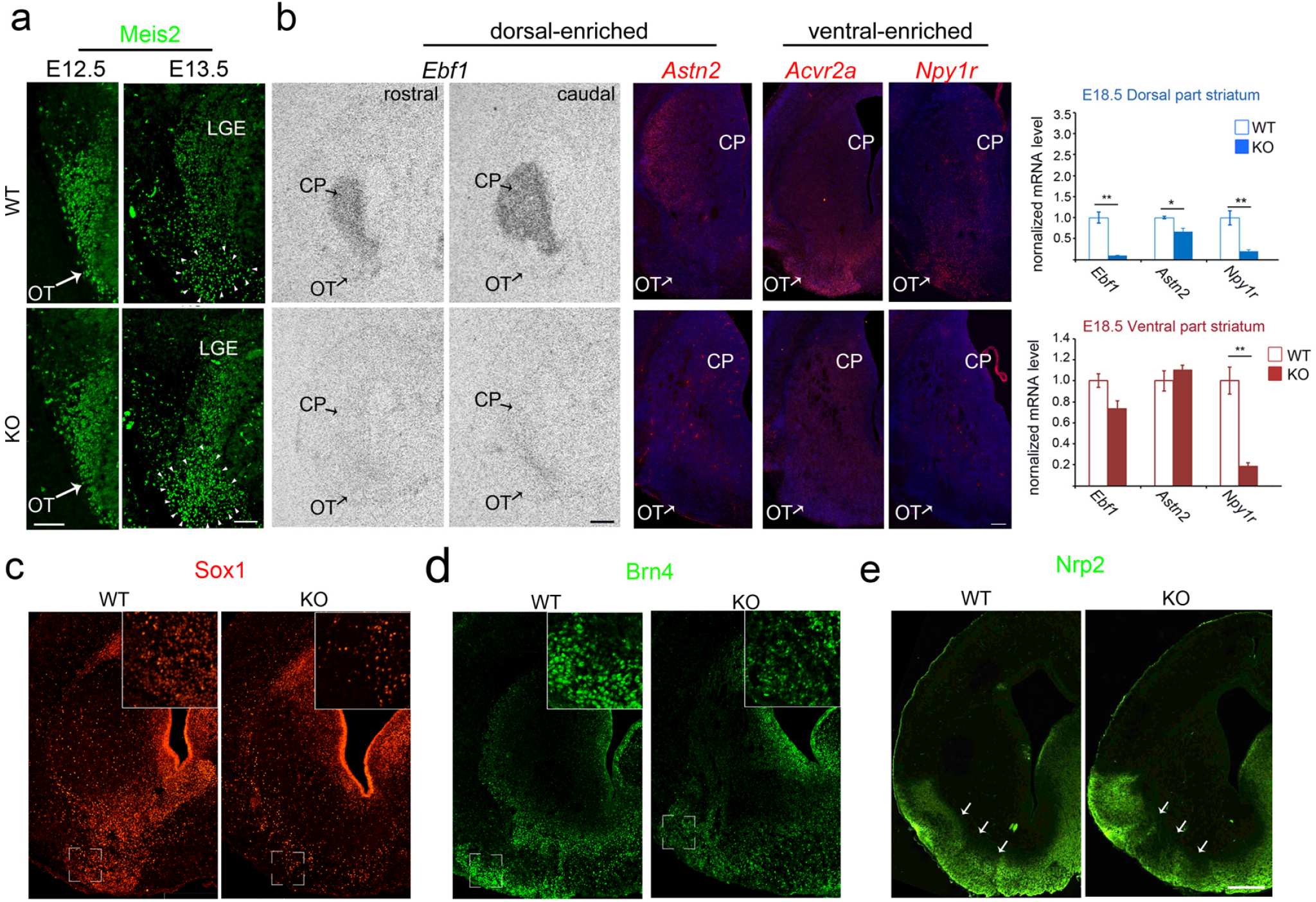
Altered gene expression patterns in the ventral striatum of Nolz-1 knockout brains. **(a)** No apparent change of Meis2 immunostaining is detected in E12.5 striatal anlage. By E13.5, increased Meis2 immunostaining was found in the ventral striatal anlage. **(b)** The ventral striatum-enriched genes *Acvr2a* and *Npy1r* are decreased in the ventral striatum of E18.5 Nolz-1 KO brains. The dorsal striatum-enriched genes, *Ebf1* and *Astn2*, remain low levels in the ventral KO striatum, though *Ebf1* and *Astn2* are decreased in dorsal KO striatum. The qRT-PCR confirmed the altered patterns of *Ebf1*, *Astn2* and *Npy1r* expressions in dorsal and ventral striatum of E18.5 Nolz-1 KO brains. **(c, d)** Sox1 **(c)** and Brn4 **(d)** immunostaining are reduced in the ventral striatum of E18.5 Nolz-1 knockout brains. **(e)** In wild type brains. Neuropilin 2 (Nrp2) is expressed from the septum across the olfactory turbercle (OT) to the piriform cortex. In Nolz-1 KO brains, an indented pattern (arrows) of Nrp2 immunostaining was found in the ventral striatum. *, p < 0.05; **, p < 0.01, n = 3/group. Scale bars in **(a)**, 100 μm, in **(b)**, 500 μm; in **(e)** for **(c-e)**, 200 μm.

**Supplementary Figure 4.**
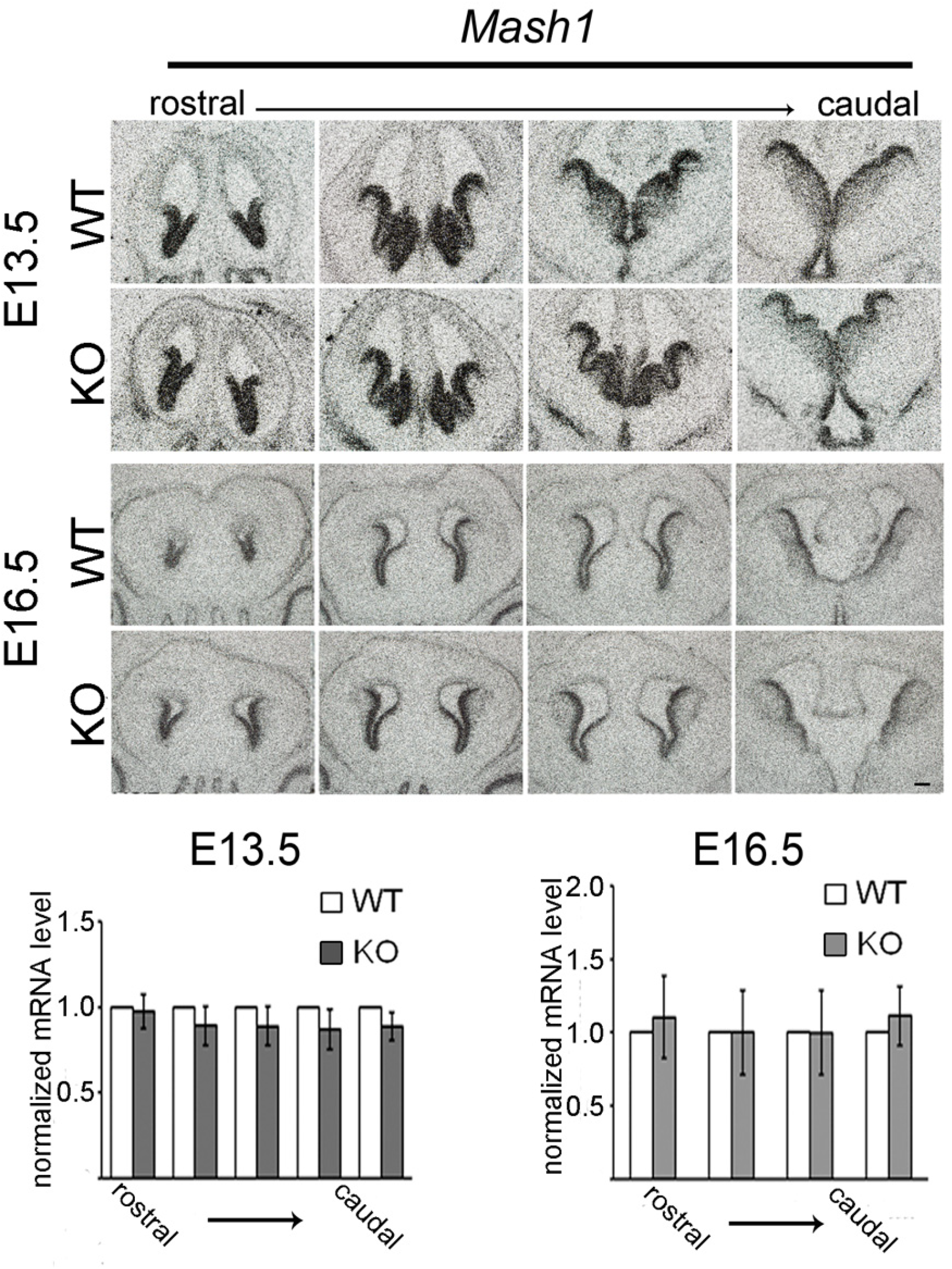
The expression of *Ascl1/Mash1* proneural gene is not altered in *Nolz-1* knockout striatal anlage. The expression pattern of *Ascl1/Mash1* mRNA is not altered in E13.5 and E16.5 from rostral to caudal levels. n = 3/group. Scale bar, 100 μm.

**Supplementary Figure 5.**
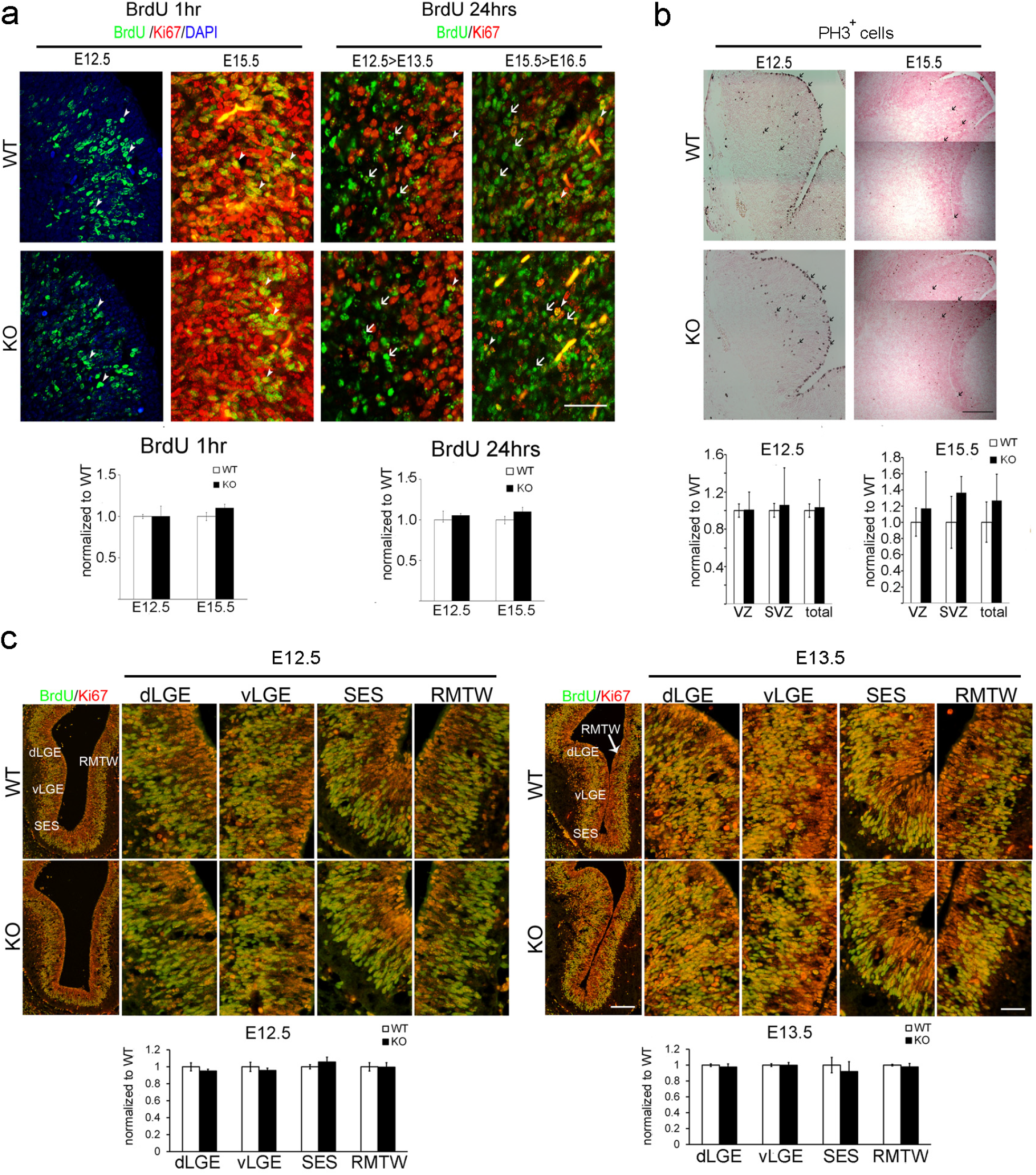
The proliferation and cell cycle exit indexes are not changed in *Nolz-1* KO striatal anlage. **(a)** BrdU was injected at E12.5 or E15.5, and the brains were collected 1 hr or 24 hr later. For BrdU labeling 1 hr after injection, the ratio of BrdU^+^;Ki67^+^ cells/total Ki67^+^ cells is used as an index for for cell proliferation. For 24 hr after BrdU injection, the ratio of BrdU^+^;Ki67^-^ cells/total BrdU^+^ cells is used as an index for the cell cycle exit index. There is no change of proliferation and cell cycle exit indexes in *Nolz-1* KO striatum. **(b)** Immunostaining of phospho-histone3 (PH3) showed that scattered PH3^+^ mitotic cells are distributed in E12.5 and E15.5 striatal anlage. No change of PH3^+^ cells were found in *Nolz-1* KO striatal anlage. **(c)** No changes in proliferation index were detected in the progenitor domains of dorsal LGE (dLGE), ventral LGE (vLGE), septoeminential sulcus (SES) and rostromedial telencephalic wall (RMTW) in E12.5 and E13.5 Nolz-1 KO forebrains.: n = 3/group. Scale bars in **(a)**, 40 μm; in **(b)**, 200 μm; in **(c)**, 100 μm and 50 μm for low and high magnification, respectively.

**Supplementary Figure 6.**
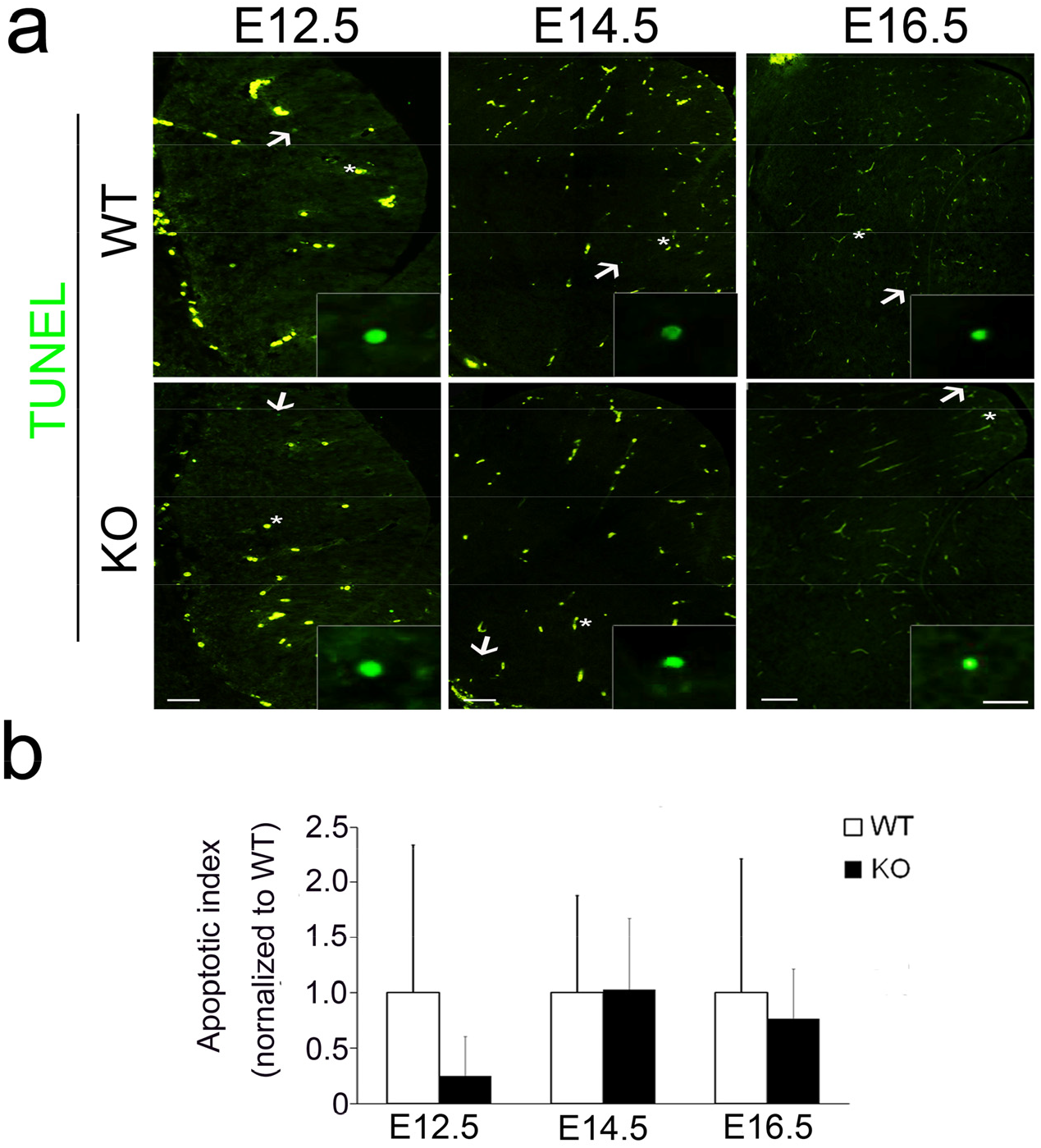
Cell apoptosis is not changed in *Nolz-1* KO striatum. **(a)** TUNEL assay is performed at E12.5, E14.5 and E16.5 wildtype (WT) and Nolz-1 knockout (KO) brains. The green round signals (insets) are counted as TUNEL^+^ apoptotic cells (arrows). Non-specific vascular staining is indicated by asterisks. **(b)** Quantification shows no change in TUNEL^+^ apoptotic cells in *Nolz-1* KO striatum. n = 3/group. Scale bars in **(a)**, 200 μm for E12.5 and E14.5; 250 μm for E16.5; 10 μm for insets.

**Supplementary Figure 7.**
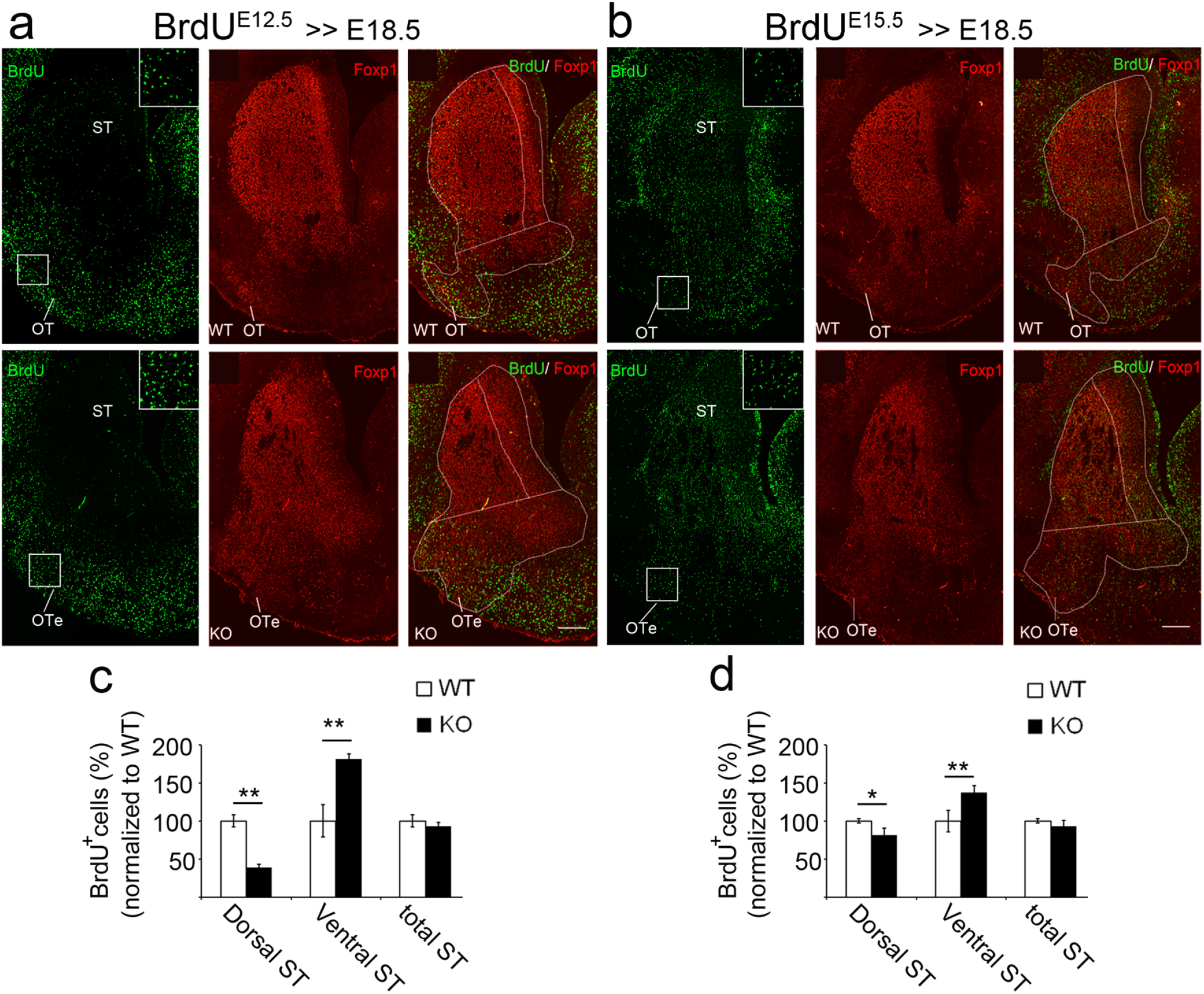
Re-distribution of BrdU^+^ cells from the dorsal to ventral striatum in *Nolz-1* KO brains. **(a, c)** BrdU was pulse-labeled at E12.5 and then analyzed at E18.5. Foxp1^+^ area was used to define the total striatal complex area (dorsal and ventral striatum). The line between the septoeminential sulcus and the piriform cortex was used to separate the dorsal from ventral striatum. The green round signals are counted as BrdU^+^ cells (insets). BrdU^E12.5^ cells were decreased in dorsal *Nolz-1* KO striatum. In contrast, BrdU^E12.5^ cells were markedly increased in the enlarged olfactory tubercle (OTe) of *Nolz-1* KO ventral striatum. **(b, d)** Similar to the distribution of BrdU^E12.5^ cells, BrdU^E15.5^ cells were decreased in dorsal striatum, but increased in ventral striatum of Nolz-1 KO striatum. The total numbers of BrdU^E12.5^ cells **(c)** or BrdU^E15.5^ cells **(d)** are not changed in Nolz-1 KO striatum. *, p < 0.05; **, p < 0.01. n = 3/group. Scale bars in **(a, b)**, 200 μm.

**Supplementary Figure 8.**
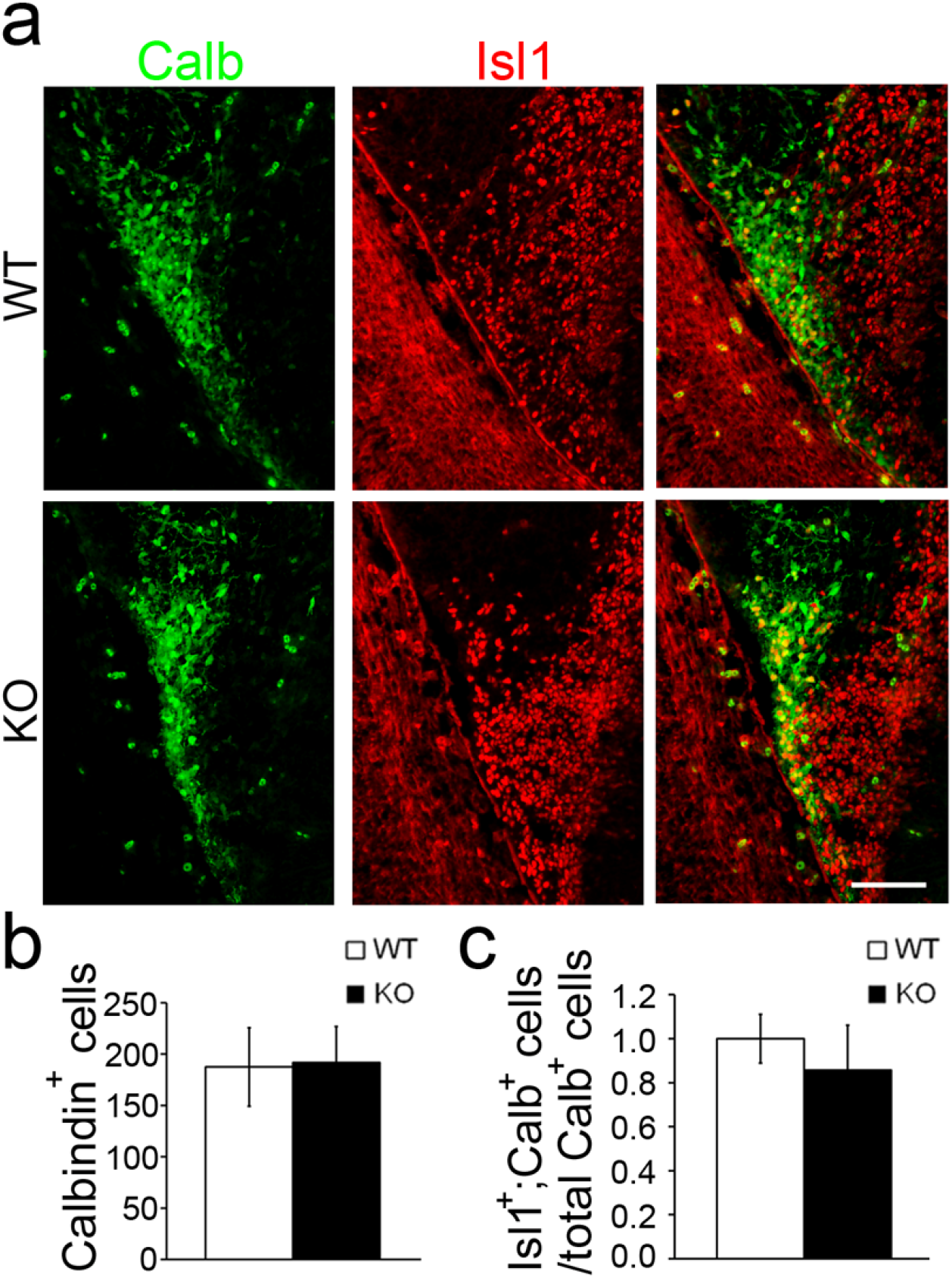
We further attempted to clarify the identify of Isl1^+^ cells in mutant OT by double immunostaining of Isl1 (red) and calbindin (green), a marker of OT neurons at early stages (Garcia-Moreno et al., 2008). **(a)** Double immunostaining of Isl1 (red) and calbindin (green), a marker of OT neurons. **(b)** Quantification shows that the number of calbindin^+^ cells is not changed in the presumptive olfactory tubercle (OT) of E13.5 Nolz-1 knockout (KO) brains. **(c)** The percentage of Isl1^+^;calbindin^+^ cells/calbindin^+^ cells is not changed in the presumptive OT of Nolz-1 KO brains. n = 3/group. Scale bar, 50 μm.

**Supplementary Figure 9.**
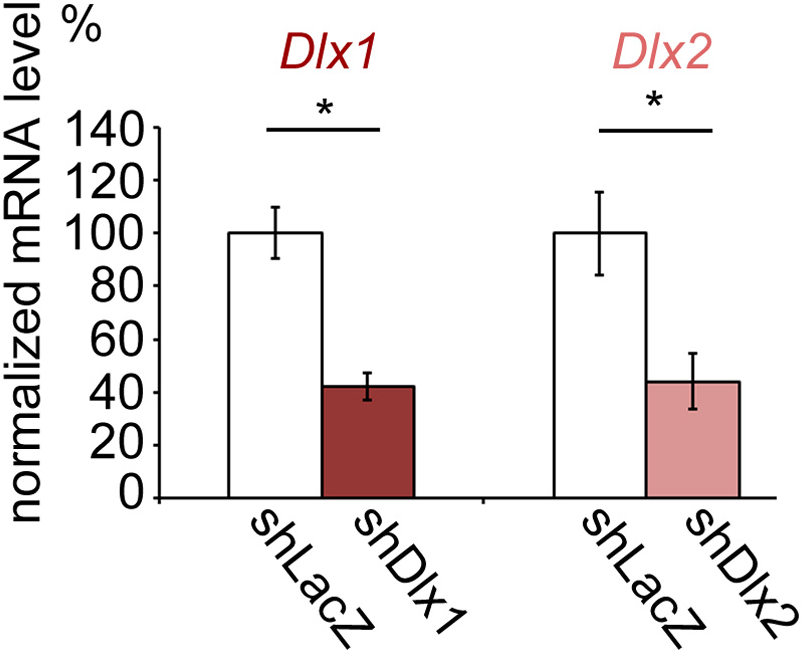
Validation of knockdown efficiency of Dlx1 and Dlx2 shRNAi. Dlx1 and Dlx2 shRNAi plasmids were transfected into P19 cells. qRT-PCR assays show that *Dlx1* and *Dlx2* mRNA levels are decreased by 58% and 56% with shDlx1 and shDlx2 RNAi, respectively. *, p < 0.05. n = 3/group.

**Supplementary Table 1.**
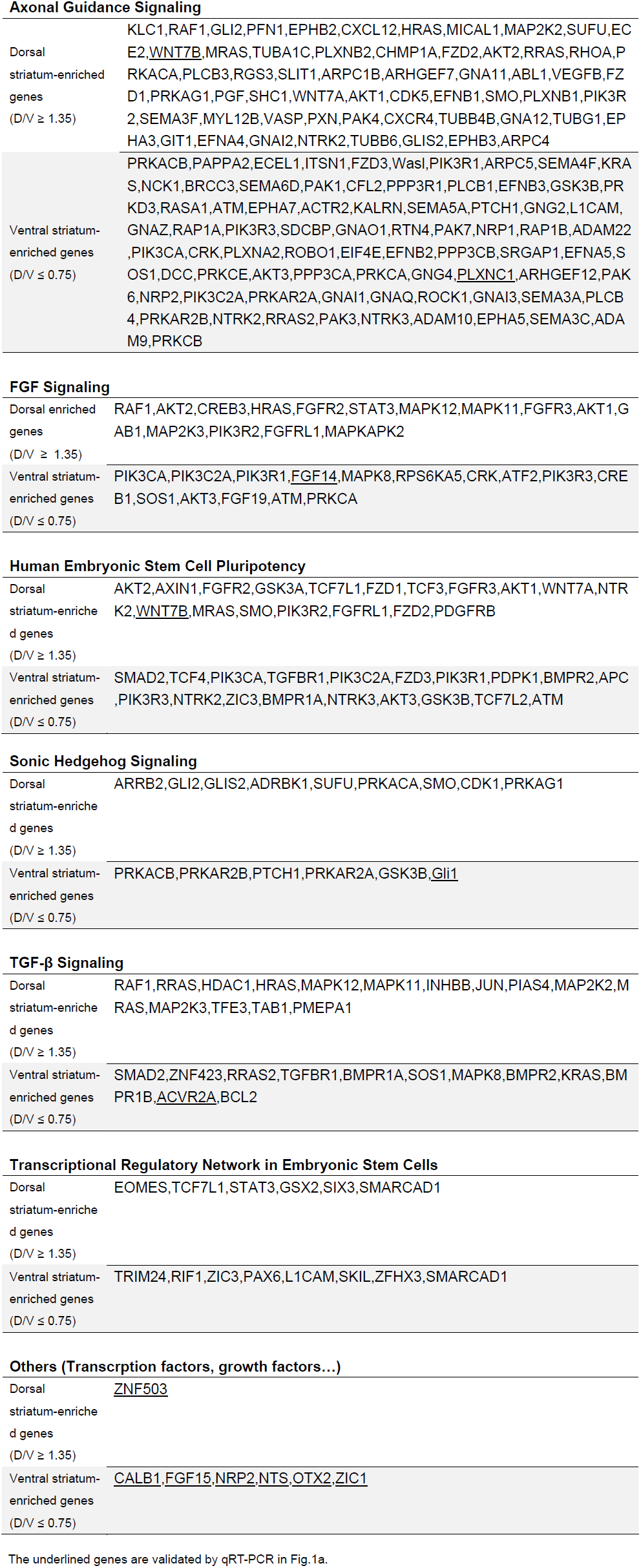
Ingenuity Pathway Analyses of dorsal or ventral striatum-enriched genes in E13.5 mouse brains.

Axonal Guidance Signaling
|  |  |
| --- | --- |
| Dorsal striatum-enriched genes<br>(D/V $\geq 1.35$) | KLC1,RAF1,GLI2,PFN1,EPHB2,CXCL12,HRAS,MICAL1,MAP2K2,SUFU,EC E2,<u>WNT7B</u>,MRAS,TUBA1C,PLXNB2,CHMP1A,FZD2,AKT2,RRAS,RHOA,PRKACA,PLCB3,RGS3,SLIT1,ARPC1B,ARHGEF7,GNA11,ABL1,VEGFB,FZD1,PRKAG1,PGF,SHC1,WNT7A,AKT1,CDK5,EFNB1,SMO,PLXNB1,PIK3R2,SEMA3F,MYL12B,VASP,PXN,PAK4,CXCR4,TUBB4B,GNA12,TUBG1,EPHA3,GIT1,EFNA4,GNAI2,NTRK2,TUBB6,GLIS2,EPHB3,ARPC4 |
| Ventral striatum-enriched genes<br>(D/V $\leq 0.75$) | PRKACB,PAPPA2,ECEL1,ITSN1,FZD3,Wasl,PIK3R1,ARPC5,SEMA4F,KRAS,NCK1,BRCC3,SEMA6D,PAK1,CFL2,PPP3R1,PLCB1,EFNB3,GSK3B,PRKD3,RASA1,ATM,EPHA7,ACTR2,KALRN,SEMA5A,PTCH1,GNG2,L1CAM,GNAZ,RAP1A,PIK3R3,SDCBP,GNAO1,RTN4,PAK7,NRP1,RAP1B,ADAM22,PIK3CA,CRK,PLXNA2,ROBO1,EIF4E,EFNB2,PPP3CB,SRGAP1,EFNA5,SOS1,DCC,PRKCE,AKT3,PPP3CA,PRKCA,GNG4,<u>PLXNC1</u>,ARHGEF12,PAK6,NRP2,PIK3C2A,PRKAR2A,GNAI1,GNAQ,ROCK1,GNAI3,SEMA3A,PLCB4,PRKAR2B,NTRK2,RRAS2,PAK3,NTRK3,ADAM10,EPHA5,SEMA3C,ADAM9,PRKCB |
| FGF Signaling |  |
| Dorsal enriched genes<br>(D/V $\geq 1.35$) | RAF1,AKT2,CREB3,HRAS,FGFR2,STAT3,MAPK12,MAPK11,FGFR3,AKT1,GAB1,MAP2K3,PIK3R2,FGFRL1,MAPKAPK2 |
| Ventral striatum-enriched genes<br>(D/V $\leq 0.75$) | PIK3CA,PIK3C2A,PIK3R1,<u>FGF14</u>,MAPK8,RPS6KA5,CRK,ATF2,PIK3R3,CREB1,SOS1,AKT3,FGF19,ATM,PRKCA |
| Human Embryonic Stem Cell Pluripotency |  |
| Dorsal striatum-enriched genes<br>(D/V $\geq 1.35$) | AKT2,AXIN1,FGFR2,GSK3A,TCF7L1,FZD1,TCF3,FGFR3,AKT1,WNT7A,NTRK2,<u>WNT7B</u>,MRAS,SMO,PIK3R2,FGFRL1,FZD2,PDGFRB |
| Ventral striatum-enriched genes<br>(D/V $\leq 0.75$) | SMAD2,TCF4,PIK3CA,TGFBR1,PIK3C2A,FZD3,PIK3R1,PDPK1,BMPR2,APC,PIK3R3,NTRK2,ZIC3,BMPR1A,NTRK3,AKT3,GSK3B,TCF7L2,ATM |

